# Functionally distinct promoter classes initiate transcription via different mechanisms reflected in focused versus dispersed initiation patterns

**DOI:** 10.1101/2022.09.30.509643

**Authors:** Leonid Serebreni, Lisa-Marie Pleyer, Vanja Haberle, Oliver Hendy, Anna Vlasova, Vincent Loubiere, Filip Nemčko, Katharina Bergauer, Elisabeth Roitinger, Karl Mechtler, Alexander Stark

**Affiliations:** Research Institute of Molecular Pathology (IMP), Vienna BioCenter (VBC), Campus-Vienna-Biocenter 1, Vienna, Austria.; Institute of Molecular Biotechnology (IMBA), Vienna BioCenter (VBC), Dr. Bohr-Gasse 3, Vienna, Austria.; Medical University of Vienna, Vienna BioCenter (VBC), Vienna, Austria.

## Abstract

Recruitment of RNA polymerase II (Pol II) to promoter regions is essential for transcription. Despite conflicting evidence, the Pol II Pre-Initiation Complex (PIC) is often thought to be of uniform composition and assemble at all promoters via an identical mechanism. Here, we show using *Drosophila melanogaster* S2 cells as a model that promoter classes with distinct functions and initiation patterns function via PICs that display different compositions and dependencies: developmental promoter DNA readily associates with the canonical Pol II PIC, whereas housekeeping promoter DNA does not and instead recruit different factors such as DREF. Consistently, TBP and DREF are required by distinct sets of promoters, and TBP and its paralog TRF2 function at different promoter types, partly exclusively and partly redundantly. In contrast, TFIIA is required for transcription from all promoters, and we identify factors that can recruit and/or stabilize TFIIA at housekeeping promoters and activate transcription. We show that promoter activation by these factors is sufficient to induce the dispersed transcription initiation patterns characteristic of housekeeping promoters. Thus, different promoter classes direct distinct mechanisms of transcription initiation, which relate to different focused versus dispersed initiation patterns.

## Introduction

Transcription of protein-coding genes by RNA polymerase II (Pol II) is a highly regulated process orchestrated by non-coding regulatory elements, namely enhancers and promoters. Pol II recruitment at promoters leads to transcription initiation from the core-promoter region, a roughly 100 base-pair region around the transcription start site (TSS) at the 5’end of protein-coding genes (Butler and Kadonaga, 2002). Although core-promoter DNA fragments on their own are typically not sufficient for activity *in vivo* and support only low levels of transcription *in vitro* (Juven-Gershon & Kadonaga, 2010), the TATA-box core-promoter is sufficient to bind the TATA-binding protein (TBP) and assemble the Pol II preinitiation complex (PIC) (Buratowski et al. 1989; Geiger et al. 1996; Petrenko et al. 2019; see also below). This finding suggests that the core-promoter DNA sequence has a crucially important function for PIC assembly and transcription, and made the TATA-box core promoter subtype a prominent model for studies of PIC assembly and transcription initiation (Smale and Kadonaga, 2003).

There are two broad classes of promoters in *Drosophila melanogaster*: 1) developmental promoters contain TATA-boxes, downstream promoter elements (DPEs), and/or Initiator (INR) motifs (Carninci et al., 2006; Ohler et al., 2002; Vo Ngoc et al., 2017; 2020), whereas 2) housekeeping promoters contain TCT, DRE and Ohler1/6 motifs (Figure 1A). These two classes of promoters exhibit distinctive regulatory properties, respond differently towards activating cues (Arnold et al., 2016; Zabidi et al., 2015), and are activated by distinct sets of coactivators (Haberle et al., 2019). In addition, developmental promoters typically display focused initiation at a single, dominant TSS, whereas housekeeping promoters typically display dispersed initiation at multiple TSSs (Rach et al., 2011).

**Figure 1.**
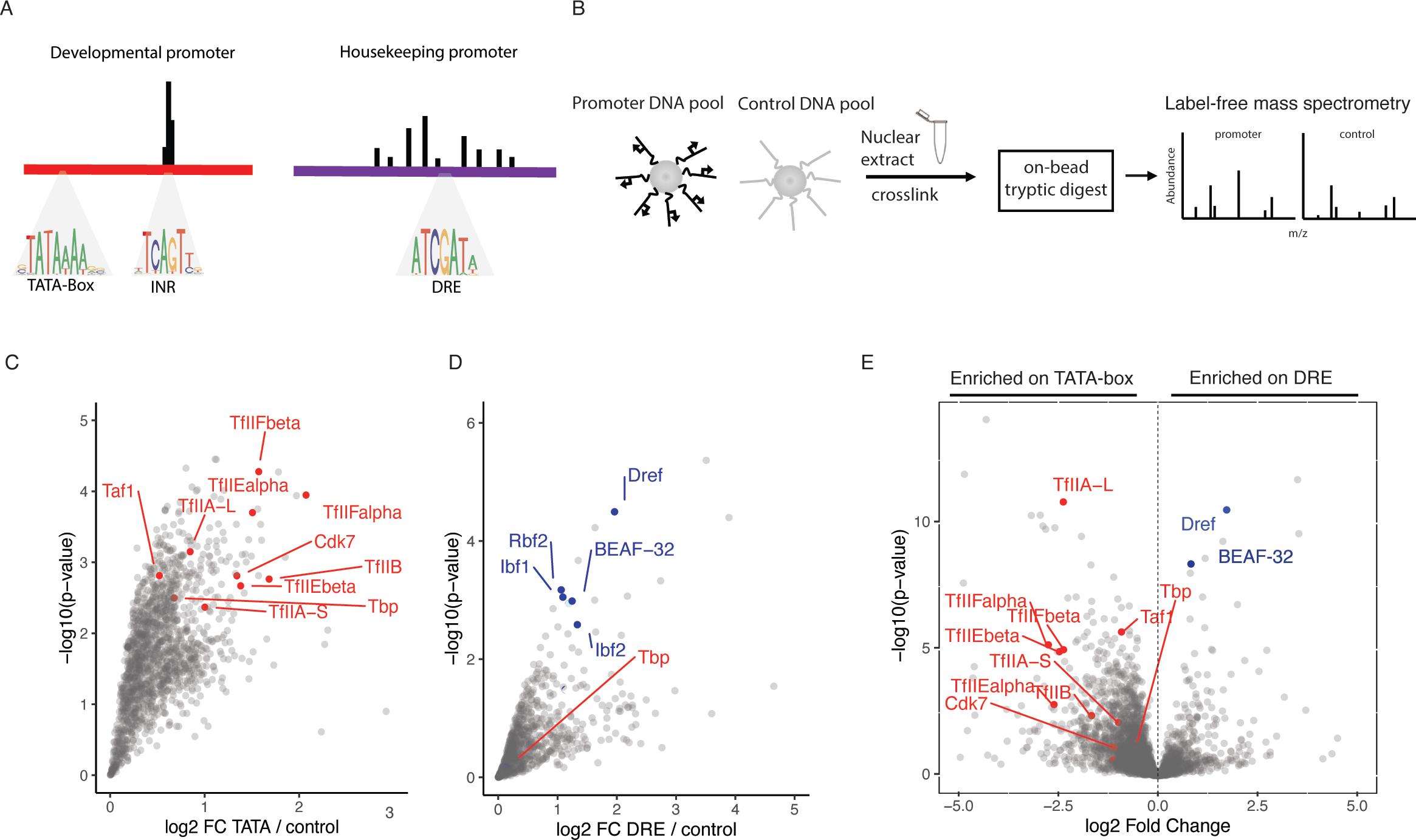
DNA affinity purifications uncover differentially bound proteins at functionally distinct promoters. A. Examples of stereotypical 121bp long core-promoters used for DNA-affinity purification of a developmental TATA-box type, and a DRE motif containing housekeeping type. TATA-box promoters exhibit focused transcription initiation from the INR motifs focused around 1-3bp, while DRE containing promoters exhibit disepersed transcription initiation across 50-100bp. B. Scheme of DNA affinity purification coupled to label free mass spectrometry. Promoters are analyzed in pools and enrichment is measured against a pool of negative control regions from the *Drosophila* genome. C. Enrichment of proteins detected by mass spectrometry on a pool of TATA-box promoters over control DNA sequences, and in D. over a pool of DRE promoters. 3 biological replicates were performed for each promtoer and control pool, significance measured with a Limma p-value<0.05. E. Enrichment of proteins bound to DRE promoters over TATA-box promoters. Limma p-value<0.05.

The general transcription factors (GTFs: TFIIA, TFIIB, TFIID, TFIIE, TFIIF, and TFIIH) assemble the PIC hierarchically at TATA-box core-promoters: the TATA-binding protein (TBP) within TFIID binds to the TATA-box motif in promoter DNA and recruits TFIIA, followed by the remaining GTFs (Cosma, 2002; Orphanides et al., 1996) and Pol II. TFIIA cooperates with TFIID to commit PIC assembly into an active state on promoters *in vitro* (Buratowski et al., 1989; Papai et al., 2010; Warfield et al., 2017). However, the nature of the PIC and PIC assembly at different core-promoter subtypes and whether they relate to these promoters’ distinct functions, remain unknown; moreover, the distinct properties of core promoter subtypes seem incompatible with a single mechanism of PIC assembly and transcription initiation.

Some evidence indeed suggests that different promoters utilize different PIC components. For example, some cells don’t seem to require TBP (Wieczorek et al., 1998; Gazdag et al., 2016; Martianov et al., 2002; Kwan et al., 2021) and some promoters require only a subset of GTFs for transcription *in vitro* (Parvin et al., 1994; 1992) or in cells (Santana et al., 2022), which is in-line with the existence of different stable intermediates or alternative arrangements of the PIC on promoter DNA (Buratowski et al., 1989; Murakami et al., 2013; Wieczorek et al., 1998; Yudkovsky et al., 2000; Yu et al., 2020). Further, promoter-bound multi-subunit protein complexes that are part of the PIC, such as TFIID, can exhibit different arrangements. For instance, the Taf9 subunit of TFIID regulates cell-type-specific genes in neural stem cells (Neves and Eisenman, 2019), whereas the Taf3 subunit of TFIID activates cell-type specific genes in myoblasts (Stijf-Bultsma et al., 2015).

In addition, some GTFs might not be required in all cells (Kwan et al., 2021; Martianov et al., 2002; Gazdag et al., 2016; Cabart et al., 2011) and/or GTF paralogs may regulate transcription in distinct cell types or at specific promoters (Akhtar and Veenstra, 2011; Duttke et al., 2014; Zehavi et al., 2015). The TBP-related factors TBP2 (also known as TRF3) and TBPL1 (TRF2 in Drosophila) have for example been implicated in transcription in early steps of mouse oocyte differentiation and during spermatogenesis, respectively (Martianov et al., 2016; Zhang et al., 2001; Yu et al., 2020; Gazdag et al., 2009), In *Drosophila*, Trf2 has been suggested to regulate the transcription of ribosomal protein genes, histone H1, and DPE motif-containing promoters (Baumann and Gilmour, 2017; Isogai et al., 2007; Kedmi et al., 2020; Wang et al., 2014). This cumulative evidence suggests that different promoter-bound GTF assemblies may exist on different promoter types and/or in different cell types, which potentially relates to these promoters’ distinct properties.

Here, we used DNA-affinity purification to identify proteins that closely interact with core-promoters, combined with protein depletion and PRO-seq to identify proteins that are required for the transcriptional function of core-promoters. We found differential use of TBP and Trf2 at different promoter subtypes and discovered distinct recruitment mechanisms of TFIIA: TFIIA was enriched at developmental promoters *in vitro* and required for their activity *in vivo*, suggesting a direct recruitment mechanism and compact PIC architecture at this promoter class. In contrast, TFIIA was not enriched at housekeeping promoters *in vitro* but still required for their activity *in vivo*, suggesting an indirect recruitment mechanism and/or dispersed PIC architecture at these promoters. Our work suggests that direct recruitment of TFIIA at developmental promoters leads to their focused initiation pattern, whereas indirect recruitment of TFIIA at housekeeping promoters leads to their dispersed initiation pattern.

## Results

### In vitro DNA-affinity purification detects core-promoter DNA – protein interactions

Roughly 37% of core promoters in the Drosophila genome can be classified as developmental (TATA+INR, DPE+INR, INR only), and 38% as housekeeping (Ohler1/6, DRE, TCT), based on previous work by others and us (Supplementary Figure 1A) (Haberle and Stark, 2018; Ohler et al., 2002; Vo Ngoc et al., 2019). Given the distinct sequences and regulatory functions of these two types of core promoters, we hypothesized that the core-promoter DNA directly binds to different transcription-related proteins. Using TATA-box core-promoters as positive control and reference point, we reasoned that short (121 bp) core-promoter DNA fragments of the different core-promoter types might bind different sets of proteins and that these could be identified *in vitro*, using conditions that assemble the canonical PIC on TATA-box promoters *in vitro* (Geiger et al., 1996; Nikolov et al., 1995; Plaschka et al., 2015; Tan et al., 1996). We therefore selected core-promoter fragments that are not themselves transcriptionally active, yet are readily inducible by activators to drive high levels of transcription in luciferase assays (Supplementary Figure 2B and 2C).

**Figure 2.**
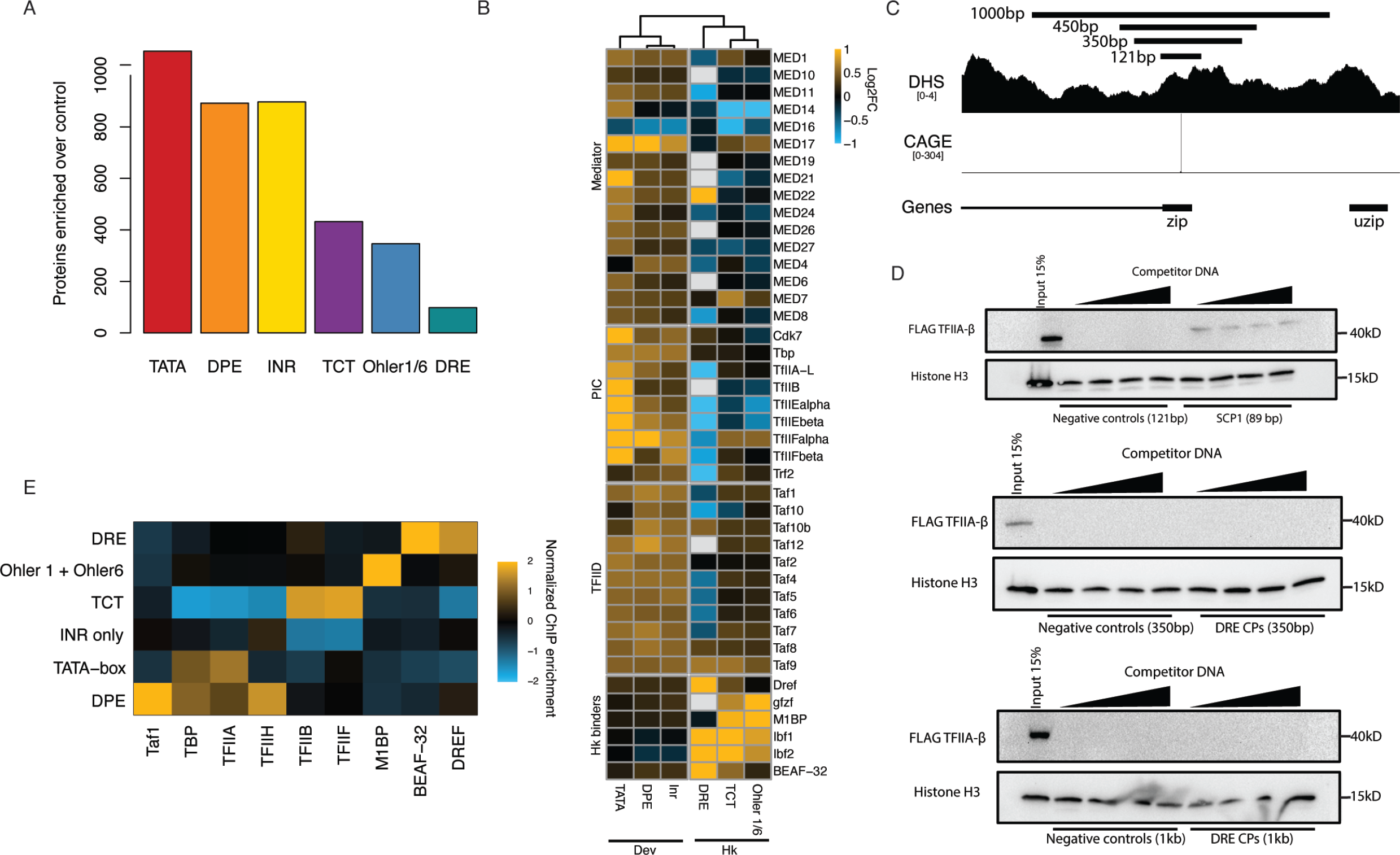
Developmental and housekeeping promoters bind different sets of proteins and GTFs. A. Total number of enriched proteins on the different tested pooled promoter types from the DNA affinity purification mass spectrometry (Limma p-value<0.05 enrichment>0). B. Enrichments from DNA affinity purification mass spectrometry of selected proteins and protein complexes on the tested different pooled promoter types compared to negative control DNA. White represents protein not detected in the given sample. Three biological replicates per condition with a Limma p-value<0.05. C. Tested regions around the zip promoter that were used in DNA-affinity purification and luciferase assay around the zip gene promoter. CAGE and DHS indicate that this promoter is accessible and transcribed in S2 cells. D. Western blots of the DNA-bound fraction eluted off the beads in a DNA-affinity purification assay. Super core promoter 1 (SCP1) was used a positive control to bind TFIIA-β-FLAG (top panel). DRE promoter pools of varying lengths were assayed for their ability to bind TFIIA. Sonicated salmon sperm DNA was used as competitor DNA and titrated from 100ng to 1.6ug per reaction. Histone H3 was used as an abundant non-specific DNA interacting protein for loading control. E. ChIP-seq signal of GTFs and select housekeeping promoter binders from Drosophila S2 cells and embryos normalized to nascent transcription level as measured by PRO-seq at the respective promoter types.

First, we examined TATA-box-containing developmental core promoters and DRE-containing housekeeping core promoter subtypes. To detect proteins that directly bind different promoter sequences of the same subtype, we pooled 16-32 representative core promoters per subtype, and used a pool of 18 non-promoter control DNA fragments as a negative control (Figure 1B). We coupled the fragments of each pool to streptavidin-coated beads, incubated the beads with S2 cell nuclear extract and free competitor DNA, washed and cross-linked associated proteins, and quantified the enriched proteins by label-free mass spectrometry (Figure 1B). We performed three replicate experiments per pool and detected between 30-35 thousand peptides each, which allowed the label-free quantification of 3,465 proteins in total across all samples. Using the three replicates, we detected 1094 proteins significantly enriched at the TATA-box core promoters over the control pool; and 98 proteins significantly enriched at the DRE core promoters (enrichment p-value < 0.05; limma (Ritchie et al., 2015)).

As expected from previous biochemical and structural work (Geiger et al., 1996; Nikolov et al., 1995; Plaschka et al., 2015; Tan et al., 1996), the TATA-box containing core promoters were enriched for the canonical Pol II PIC, including TBP, GTFs and TFIID, and most Mediator subunits (Figure 1C, and Supplementary Figure 1B), confirming that TATA-box promoter DNA is sufficient to directly bind these proteins *in vitro* and that our setup captures these protein-DNA complexes.

Unexpectedly, the DRE-containing core promoters did not enrich for any of the Pol II PIC subunits; indeed some Tafs and GTFs were even depleted compared to control DNA. In contrast, the DRE core promoters were enriched for the core-promoter-element binding factor DREF, BEAF-32 and Ibf1/2 among other proteins (Figure 1D). Directly plotting the enrichments at DRE versus TATA promoters confirmed the strong differential recruitment of GTFs and PIC components specifically to TATA promoters but not to DRE promoters (Figure 1E). Mutating either the TATA-box or DRE motifs reduced TBP and DREF binding to control levels, respectively (Supplementary Figure 1B), suggesting that the differential binding of these proteins is directed by the different promoter DNA sequences as expected (Tora and Timmers, 2010; Kwon et al., 2003).

### Different promoter subclasses show distinct binding of the Pol II PIC

Our DNA-affinity purification detected an association between known PIC components and TATA-box-containing developmental core-promoters, but not with housekeeping DRE core-promoters. To determine if the results above generalize to other promoter subtypes, we extended our analysis to additional developmental promoters containing DPE or INR motifs, and to housekeeping promoters containing TCT or Ohler 1/6 motifs.

We found that developmental promoter subtypes enriched for 892 to 1093 proteins, whereas housekeeping promoter subtypes enriched only between 98 and 432 proteins (enrichment p-value<0.05) (Figure 2A, Supplementary Table 1). Moreover, developmental and housekeeping promoters enriched for different sets of proteins: GTFs and PIC components were preferentially enriched at all developmental promoters but were not or only weakly enriched at housekeeping promoters (Figure 2B). Similarly, multiple components of the Mediator and TFIID complexes were preferentially enriched at developmental promoters, with TATA-box containing promoters showing the highest levels of binding (Figure 2B). In contrast, none of the housekeeping promoter subtypes were enriched for GTFs, TFIID or Mediator subunits; instead, they were enriched for various TFs that bind core-promoter elements and chromatin regulators. For example, DRE-containing promoters exhibited the highest enrichment of DREF and BEAF-32, whereas Ohler 1/6 promoters exhibited the highest enrichment of the Motif 1 Binding protein (M1BP) and the cofactor GFZF (Figure 2B). Our DNA-affinity purification data suggest that short DNA fragments corresponding to functionally distinct core promoters directly associate with distinct transcription-related proteins under identical conditions *in vitro*.

We considered that 121 bp was not sufficiently long for the housekeeping core-promoters to associate with the canonical PIC by DNA-affinity purification. We thus tested 350, 450 and 1000 bp long fragments derived from DRE promoters, which all bound DREF and were active even in the absence of enhancers or tethered activators (Supplementary Figure 2A and 2D). However, DNA-affinity purification still did not detect an interaction between the PIC component TFIIA-β and the longer DRE promoters which are known to bind DREF, in contrast to the TATA-box containing SCP1 promoter, a well-studied TATA-box core promoter used as a positive control (Figure 2C and 2D). Overall, DNA-affinity purification detected different sets of proteins that directly associate with housekeeping and developmental core-promoters under identical conditions *in vitro*. These findings are intriguing and suggest that the promoters’ functional differences might arise at the level of GTF recruitment and PIC assembly, presumably via distinct DNA-binding factors and/or tighter versus looser protein-DNA complex architectures.

The DNA-affinity purifications directly report the biochemical properties of the DNA fragments and suggest that core-promoter DNA fragments differ in their ability to directly bind GTFs and the PIC *in vitro*. *In vivo*, additional players, such as chromatin or nearby enhancers, can influence GTF- or Pol II recruitment and transcription initiation at core-promoters. We re-analyzed published ChIP-seq and ChIP-nexus data from Drosophila cells or embryos, which confirmed that all the assayed GTFs do indeed bind to all promoters, including housekeeping promoters (Supplementary Figure 2H & 2I).

Importantly, the ChIP signals reflected the differential binding preferences observed *in vitro* for the respective promoter subtypes (Figure 2E) (Baumann and Gilmour, 2017; Liang et al., 2014; Shao and Zeitlinger, 2017): GTFs were generally more highly enriched at developmental promoters than housekeeping promoters (except for TFIIB, TFIIF that bound strongly to TCT promoters), whereas TFs were more highly enriched at housekeeping promoters according to their motif contents: M1BP showed the highest ChIP-seq signals at Ohler 1/6 promoters, and DREF and BEAF-32 showed highest signals at DRE promoters (Figure 2E & Supplementary Figure 2E).

We infer that the DNA sequence of developmental core promoters forms a close/tight physical association with the PIC that can be detected by DNA-affinity purification. In contrast, the weaker ChIP signals and lack of DNA-affinity purification suggest a weaker/looser, less rigid, more transient, or more indirect physical association between housekeeping core-promoter DNA and GTFs. Instead, housekeeping core promoters appear to form close physical associations with sequence-specific TFs through their cognate DNA binding motifs both *in vitro* and *in vivo*. Additionally, the markedly lower number of proteins enriched at housekeeping promoters suggests that their DNA-protein interface is generally weaker, more indirect, and/or transient nature.

### Differentially recruited factors in vitro have distinct functional requirements

To determine if the differential recruitment of promoter-associated factors *in vitro* reflects distinct functional requirements *in vivo*, we used the auxin inducible degron (AID) system (Nishimura et al., 2009) to deplete endogenously labeled proteins from *D. melanogaster* S2 cells and measured nascent transcription by PRO-seq (Kwak et al., 2013), a strategy recently used for GTFs in human cells (Santana et al., 2022) (Fig. 3A).

**Figure 3.**
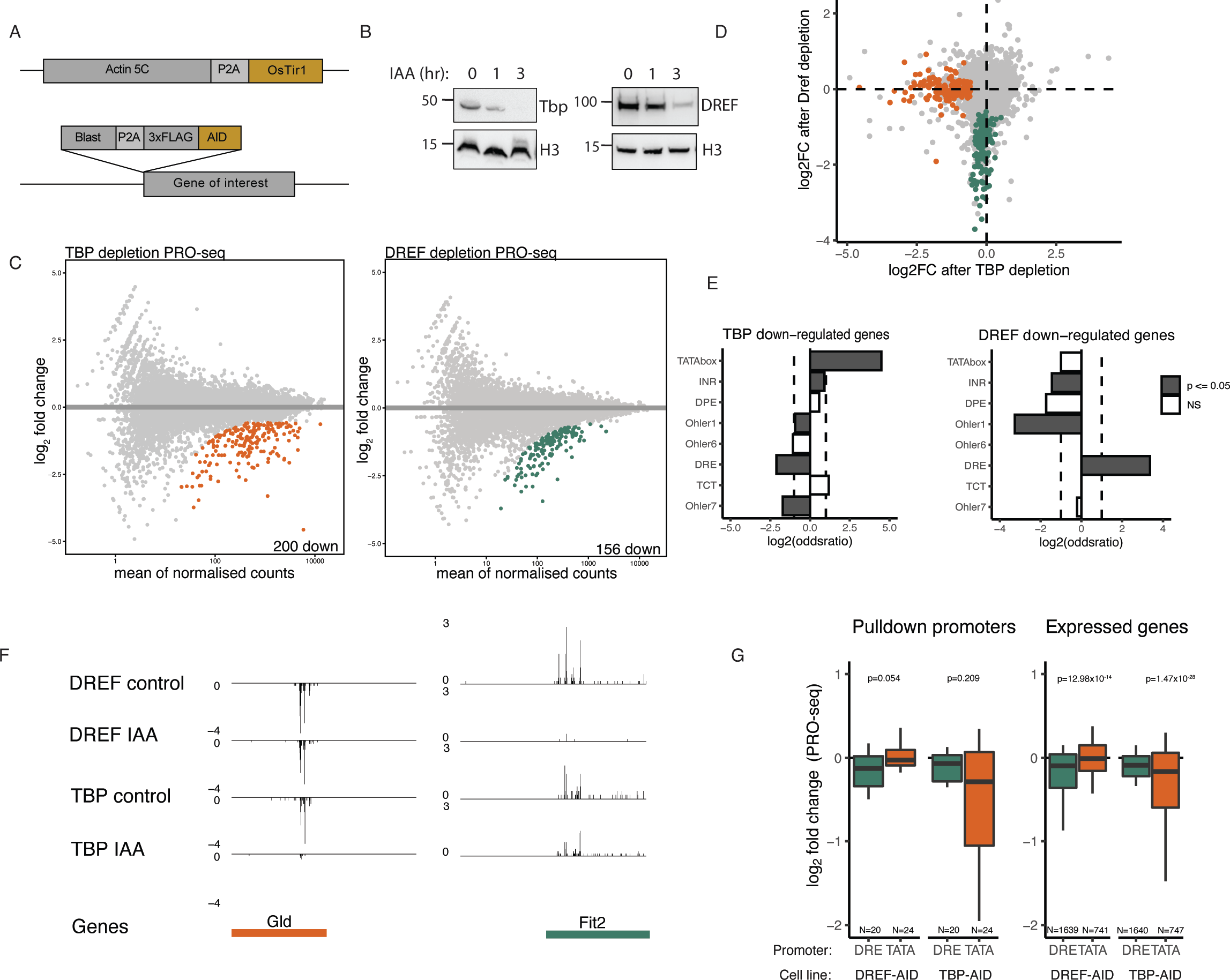
TBP and DREF are required by distinct sets of promoters. A. Strategy for generating endogenously tagged AID cell lines. An AID-3xFLAG endogenous knock-in was generated in the N-terminus of either DREF or TBP in a background cell line stably expressing the Tir1 ligase downstream of Actin5c. B. Western blot on FLAG tagged TBP and DREF 0,1 and 3 hours after auxin addition showing protein degradation C. PRO-seq measurement after 6 hours of auxin addition to the TBP or DREF AID tagged cell lines, MA plots represent colored genes which are significantly downregulated compared to no auxin control (FDR<0.05). Two biological replicates per conditions. D. Overlap of the TBP and DREF depletion PRO-seq. Green and orange colored dots represent TBP and DREF dependent promoters, fold change>-1.5 & FDR<0.05. E. Fisher’s exact test for motif enrichment in TBP and DREF downregulated promoters compared to all expressed promoters. Log2 Odds ratio displayed. F. Genome tracks of PRO-seq data indicating examples of genes that are dependent on TBP or DREF. G. Differential PRO-seq signal across TATA-box and DRE promoters which were used for the DNA affinity purification (left) or all expressed TATA-box and DREF containing promoters (right). P-values from a two-sided Wilcoxon test are provided (note that despite similar magnitude of change, the comparisons on the left are not significant due to low numbers of promoters in the compared groups.

We examined TBP and DREF first and observed the near complete degradation of both proteins three hours after auxin addition (Fig. 3B) and their complete depletion six hours after auxin addition (Supplementary Figure 3A). To ensure complete protein degradation while avoiding potential secondary effects from prolonged protein depletion, we measured changes to Pol II nascent transcription six hours after auxin treatment.

We performed two biological replicates of PRO-seq that were highly similar (PCC>0.99 Supplementary Figure 3B) and revealed 200 downregulated genes after TBP depletion and 156 downregulated genes after DREF depletion (fold-change < 0 and FDR < 0.05; Figure 3C). Notably, not a single gene was shared between the two conditions, indicating that distinct sets of promoters require TBP and DREF (Figure 3D). Motif enrichment analysis of the downregulated promoters revealed a strong enrichment of the TATA-box in the TBP-dependent promoters, and of the DRE motif in the DREF-dependent promoters (Figure 3E), as expected. The differential dependency on TBP versus DREF is apparent at the TATA-box promoter upstream of *Glucose dehydrogenase* (*Gld*) and the DRE promoter upstream of *Fermitin 2* (*Fit2*) (Figure 3F) and generalizes to the promoters used for the DNA affinity purification experiments, and to all active TATA-versus DRE-containing promoters genome-wide (Figure 3G and Supplementary Figure 3C). These results show that a relatively small number of active promoters require TBP (Martianov et al., 2002; Santana et al., 2002; Gazdag et al, 2016), and that these are specifically TATA-box containing promoters. Similarly, only a subset of promoters requires DREF, which are different from the TBP-requiring promoters and specifically contain DRE motifs. Overall, these results imply that different promoter types differentially depend on the two core-promoter element binders and utilize distinct DNA-protein interfaces and/or interactors to recruit Pol II and initiate transcription.

### TBP and TRF2 display promoter subtype-dependent requirements

As TBP seemed to be required only for TATA-box containing promoters, we wondered if TBP paralogs, specifically TRF2 (TBPL1 in mammals), might replace TBP at other promoter types (TRF, also called TRF1 is not detectable in S2 cells, Suppl. Figure 4G and 4H). In fact, TRF2 has been reported to function at DPE and TCT promoters in Drosophila (Wang et al., 2014, Zehavi et al., 2015, Kedmi et al., 2020) and we found TRF2 most strongly bound to DPE and INR containing core-promoter DNA *in vitro* (Supplementary Figure 4A; TBP bound TATA-box, DPE and INR promoters at equal levels).

**Figure 4.**
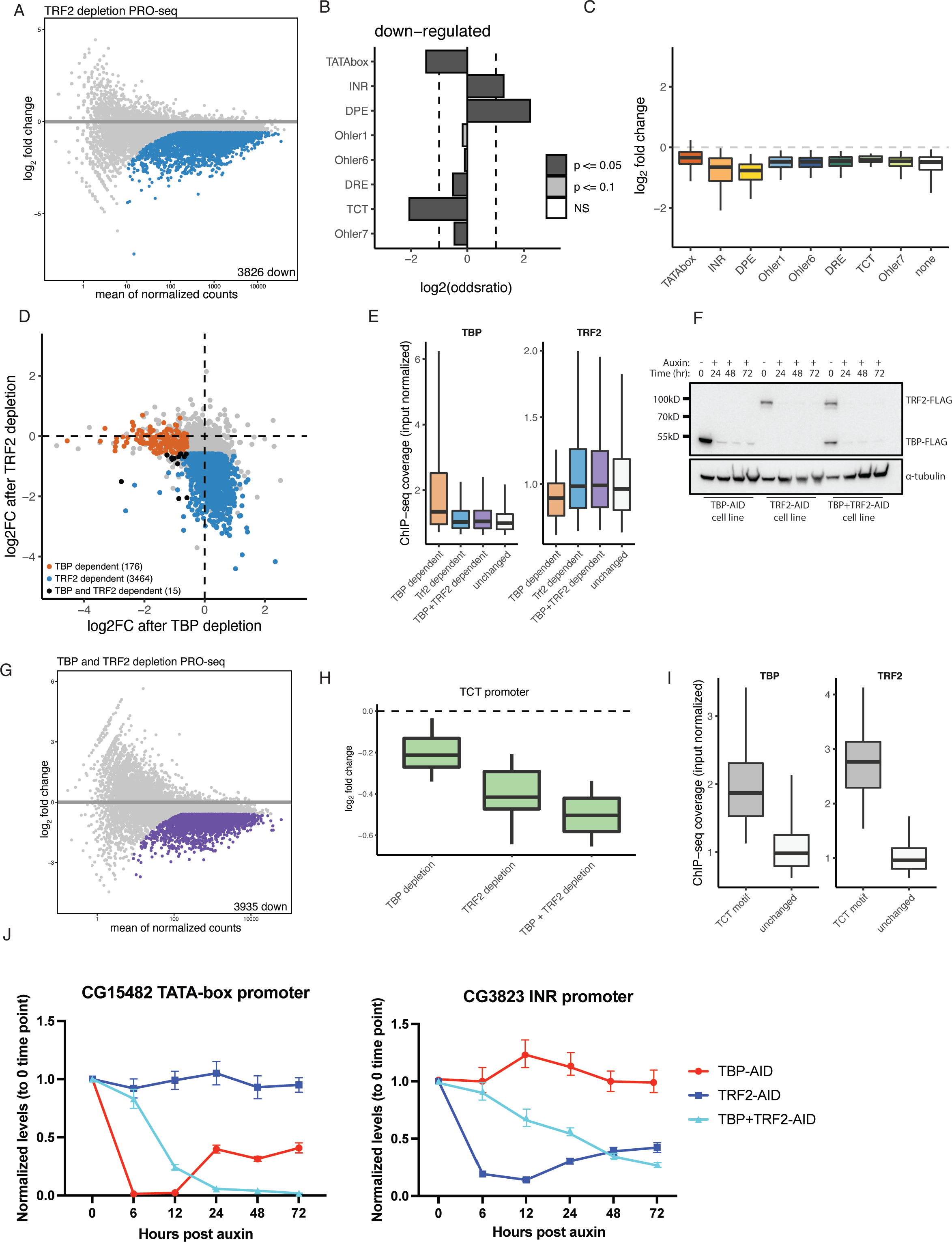
TBP and TRF2 regulate distinct subsets of developmental promoters. A. PRO-seq was performed upon a 6 hour auxin treatment of Trf2 depletion. Colored dots represent significantly downregulated genes (FDR<0.05). B. Motif enrichment analysis of gene promoters downregulated upon Trf2 depletion using a Fischer test, odds ratio is displayed. C. Box plot representation of Trf2-depletion PRO-seq data across different promoter types. Most developmental and housekeeping promoter types are affected with TATA-box promoters being least affected. D. Scatter plot of TBP and Trf2 depletion PRO-seq at 6 hours of auxin treatment. TBP dependent genes are colored in orange, Trf2 dependent genes are colored in blue. Genes dependent on both TBP and Trf2 are colored in black (FDR<0.05). E. ChIP-seq coverage (input normalized) of TBP and Trf2 at TBP or Trf2 dependent promoters, and all other active promoters labeled as ‘unchanged’. Promoters downregulated in the double depletion TBP + Trf2 cell line are colored in purple. F. Western blot of anti-FLAG antibody of TBP-AID, Trf2-AID and a double tagged TBP-AID + Trf2-AID cell lines from a multi-day time course of auxin treatment showing prolonged depletion. G. PRO-seq was performed upon a 12 hour auxin treatment of a double tagged TBP+Trf2 AID cell line. Colored dots represent significantly downregulated genes (FDR<0.05), left panel. H. PRO-seq signal of individual TBP or Trf2 and double depletion of both across TCT promoters (N=55). I. ChIP-seq coverage (input normalized) of TBP and Trf2 on TBP across TCT promoters (N=55), all other expressed by not changing promtoers are labeled as ‘unchanged’. J. qPCR on an auxin treatment time course of TBP and Trf2 dependent genes upon individual depletion of TBP or Trf2 and a double depletion of both.

To determine which promoters depend on TRF2, we AID-tagged the evolutionarily conserved short isoform of TRF2 that is expressed in S2 cells and rapidly depleted the endogenous protein by the addition of auxin. Mass spectrometric measurement of TRF2 identified peptides shared between the two isoforms, which were depleted after the addition of auxin (Supplementary Figure 4B & 4C). PRO-seq after 6 hours of auxin treatment resulted in the down regulation of 3826 genes (Figure 4A). The promoters of these TRF2-dependent genes were enriched in DPE and INR motifs, while TATA-box and TCT motifs were depleted (Figure 4B), suggesting that TBP- and TRF2-dependent genes/promoters might be different. Indeed, TRF2 depletion most strongly downregulated the INR and DPE type promoters, while TATA-box and TCT promoters were among the least affected (Figure 4C), and genes downregulated following TBP or TRF2 depletion were largely distinct (Figure 4D and Supplementary Figure 4F). Reanalysis of published ChIP-seq datasets confirms that TBP and TRF2 localize to different promoters: TBP-dependent promoters preferentially bound TBP but not TRF2 and, vice versa, TRF2-dependent promoters preferentially bound TRF2 but not TBP (Figure 4E). This mutual exclusivity suggests that DPE and INR developmental promoters and housekeeping promoters utilize TRF2 but not TBP to assemble a Pol II PIC *in vivo* at TATA-less promoters (Supplementary Figure 4K).

The depletion of TBP or TRF2 individually left approximately half of the expressed genes largely unaffected, including the TCT-promoter-bearing ribosomal protein genes, suggesting that TBP and TRF2 might function partially redundantly. We AID-tagged both genes in a single cell line (Figure 4F; see methods), which allowed the simultaneous, auxin-inducible depletion of endogenous TBP and TRF2 (albeit with slower depletion kinetics of TBP compared to TRF2 and to TBP in the TBP-AID single-tagged cell line; Supplementary Figure 4D). We performed PRO-seq after 12 hour of auxin treatment, which resulted in the downregulation not only of all three developmental promoter subtypes, but of TCT promoters (Figure 4G-I). Consistent with the downregulation of TCT promoters, the combined depletion of both TBP and TRF2 resulted in growth arrest of the auxin treated cells, starting between 24 and 48 hours after auxin treatment (Supplementary Figure 4E). The result that TCT promoters appear to function with either TBP or TRF2, which seem to function redundantly, is consistent with strong ChIP-seq signals for both TBP and TRF2 at these promoters (Figure 4H).

Surprisingly, prolonged individual depletion of either TBP or TRF2 resulted in partial recovery of transcription after 24 hours at several tested developmental promoters, however double depletion of both TBP and TRF2 resulted in continued downregulation of these genes (Figure 4J and Supplementary Figure 4H). Auxin wash-out experiments indicated that recovery of transcription of the tested genes does recover rapidly and fully (Supplementary Figure 4G). These results indicate that promoters preferentially use either TBP or TRF2 but can utilize either paralog in the absence of the other.

### All promoters – including housekeeping promoters – functionally depend on TFIIA

Our data suggest that the canonical PIC, including TFIIA, forms a closer physical association with developmental promoters when compared to housekeeping promoters. To test the functional dependency of different promoter subtypes on TFIIA, we tagged TFIIA with AID (other GTFs such as TFIIE (α and β subunit), TFIIF (α and β subunit) and TFIIB were incompatible with tagging at either the N- or C-termini and could therefore not be assessed). Given the proteolytic processing the TFIIA-L precursor protein by Taspase A to generate TFIIA-β (Yokomori et al. 1993, Zhou et al. 2006), we endogenously tagged TFIIA-L at its C-terminus, which was retained in TFIIA-β, and hereafter refer to the tagged protein as TFIIA-β and TFIIA-AID for simplicity (Figure 5A). Auxin treatment efficiently depleted TFIIA-AID within one to two hours, resulting in loss of PRO-seq signal for all expressed protein coding genes in S2 cells within 3 and 6 hours, and cell death only after 24 hours (Figure 5A-C and Supplementary 5A-D). These results suggest that TFIIA is functionally required at all promoters, including housekeeping promoters. As housekeeping promoter DNA recruits TFIIA only weakly (see above), TFIIA might be recruited to housekeeping promoters via a novel mechanism, independently of DNA-mediated recruitment of TBP.

**Figure 5.**
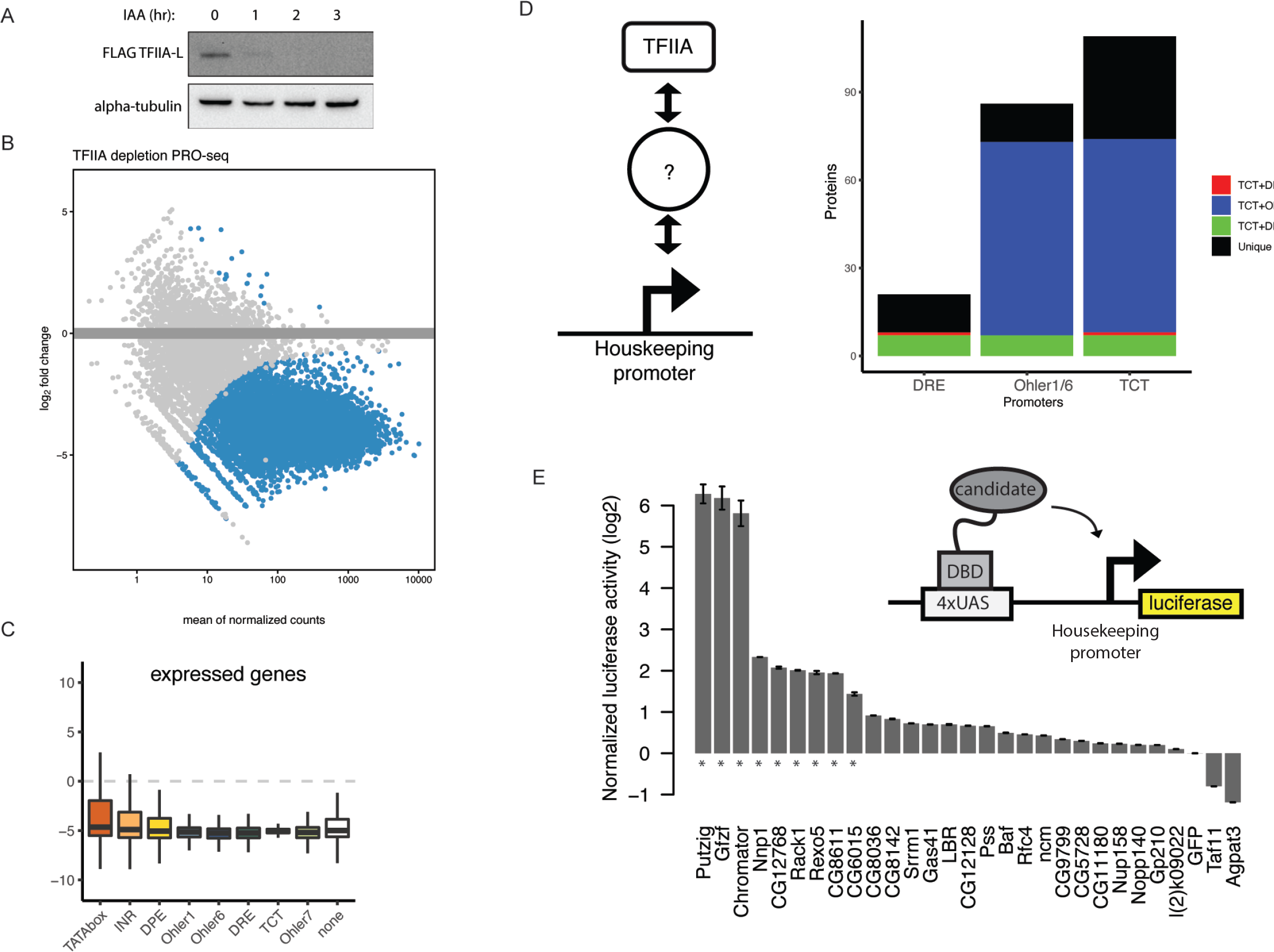
TFIIA is required by all promoters and is recruited by housekeeping cofactors to housekeeping promoters. A. Western blot for an endogenously tagged TFIIA-β-3x-FLAG-AID cell line after addition of auxin at 1, 2 and 3 hours indicating the TFIIA-β C-terminal cleaved product. B. MA-plot of PRO-seq measurement 6 hours after auxin addition to the TFIIA-β-AID cell line. Colored dots represent significant downregulation, FDR<0.05. Two biological replicates per condition. ∼10000 protein coding genes are down regulated, with 73 genes not showing down regulation due to their overlap with non-coding RNA genes such as tRNA which are not affected by TFIIA-β depletion. C. PRO-seq signal at all expressed promoters, represented according to their motif content in box plots. D. Overlap of TFIIA-β-3xFLAGimmunoprecipitation mass spectrometry data with DNA affinity purification mass spectrometry of the 3 tested housekeeping promoter types. Three biological replicates per conditions with Limma p-value<0.05 and enrichment>0. E. Luciferase assay in which Gal4 DNA binding domain fusion proteins were recruited to 4xUAS sites upstream of a minimal housekeeping Rps12 promoter. Measurements are normalized to Renilla luciferase (transfection control) and GFP. * denotes proteins activating a housekeeping promoter with a log2FC>1.5 and p-value<0.05, two-tailed student’s t-test.

### Intermediary proteins recruit TFIIA to housekeeping promoters

As housekeeping promoters depend on TFIIA for transcription in cells but fail to enrich for TFIIA by DNA-affinity purification *in vitro*, we hypothesized intermediary proteins interact with both the housekeeping promoter DNA and TFIIA to mediate PIC assembly (Figure 5D). We thus performed immunoprecipitation-mass spectrometry with the endogenously tagged TFIIA-L-AID-3xFLAG S2 cell line and the parental Tir1-expressing cell line as a control. We uncovered 300 TFIIA interacting proteins, including all three known components of the TFIIA complex and other TFIIA interactors, such as the TBP paralog TRF2 (but not TBP), members of the TFIID complex, and various GTFs, such as TFIIE (Supplementary Figure 5E and Supplementary Table 2).

To identify candidate intermediary proteins, we intersected the TFIIA binding proteins with the proteins enriched on housekeeping promoters *in vitro* (Figure 5D). Applying this strategy to developmental promoters as a positive control identified most known GTFs, thus validating the approach. We found 131 proteins that can associate with TFIIA and at least one housekeeping promoter subtype (Figure 5D), including DREF, Chromator, GFZF, Putzig, the nucleolar protein Nnp1, and the RNA helicase CG8611 (Supplementary Table 2).

To determine if the candidate TFIIA-recruiting proteins can activate transcription from a housekeeping promoter, we fused 28 candidate proteins to the Gal4 DNA-binding domain and tethered them to a UAS sequence upstream of a minimal housekeeping core promoter driving luciferase in S2 cells (Figure 5E). We found that nine proteins were able to transactivate the housekeeping promoter (fold change>4 & p<0.05), particularly the coactivators GFZF, Putzig and Chromator (Figure 5E), suggesting that they may mediate TFIIA recruitment. The top three activators: GFZF, Putzig and Chromator have previously been observed to bind housekeeping promoters, and immunoprecipitation of Chromator followed by mass spectrometry indicated these three proteins strongly interact with each other (Supplementary Figure 5F). Indeed, when we performed DNA-affinity purification with a UAS-housekeeping promoter DNA fragment, we observed co-recruitment of TFIIA with Gal4-GFZF but not Gal4-GFP onto promoter DNA *in vitro* (Supplementary Figure 5G). These data suggest that GFZF can recruit TFIIA and transactivate housekeeping promoters.

Overall, these results suggest that housekeeping promoters recruit TFIIA-β and Pol II indirectly via intermediary housekeeping cofactor proteins interacting with DNA-binding proteins, whereas developmental promoters recruit TFIIA and the PIC directly via TBP/TRF2-DNA interactions.

### Housekeeping cofactors underlie dispersed transcription initiation patterns

The results so far suggest that housekeeping promoters are unable to directly recruit a canonical PIC *in vitro* and may exhibit weaker and more indirect interactions with GTFs. We hypothesized that a less direct promoter DNA-TFIIA or DNA-PIC interface at housekeeping promoters might lead to a weak alignment between TSSs and the relevant core-promoter sequence elements, such as DREF or Ohler 1/6 motifs.

To test this hypothesis, we used Cap Analysis of Gene Expression (CAGE) data to analyze the distribution of TSSs relative to the positions of various motifs across *D. melanogster* promoters. As expected (e.g. (Ohler et al., 2002; Parry et al., 2010; Rach et al., 2011)) the TSSs of developmental promoters, such as TATA-box-, INR- or DPE-containing promoters, were restricted to a narrow window at consistent and precise distances from the core-promoter sequence elements (Figure 6A). Similarly, the TCT type housekeeping promoters exhibit a focused initiation pattern precisely at the TCT motif (Wang et al., 2014). These results confirm that initiation is precisely aligned to the TATA-box, INR, DPE and TCT motifs, as expected given previous reports and the fact that these motifs direct PIC and Pol II recruitment and initiation through TBP or TRF2 (Rach et al., 2011; Sawadogo and Roeder, 1985).

**Figure 6.**
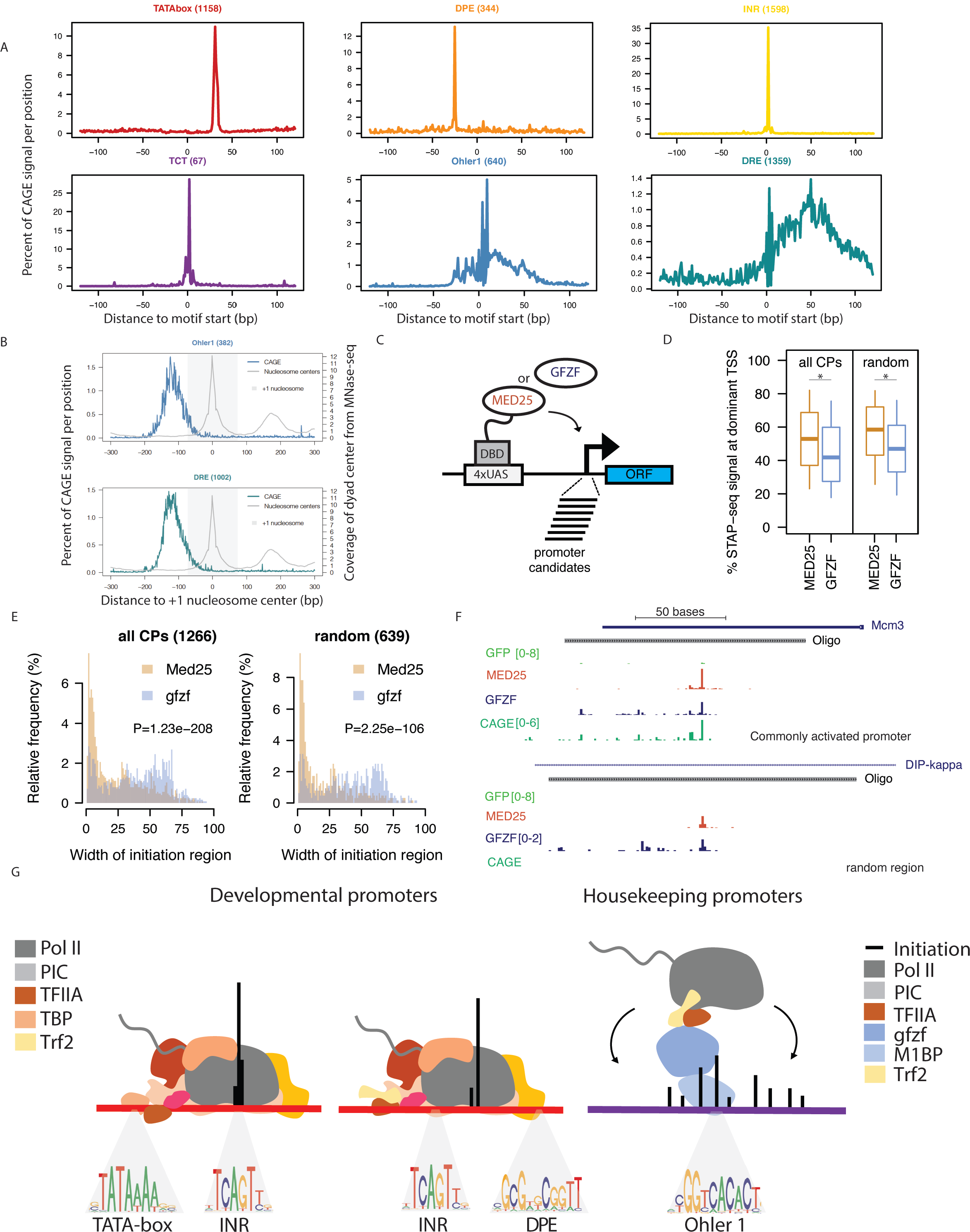
Housekeeping cofactor recruitment is sufficient to recapitulate dispersed transcription initiation patterns. A. Distribution of CAGE signal from mixed *D. mel* embryos (0-24h) centered on the location of promoter DNA motif sequence set at position 0 across the 6 main promoter types investigated in this study. B. Relative CAGE signal per position on all active promoters containing either Ohler 1 (top) or DRE (bottom) motif, aligned to the +1 nucleosome centre (point of highest coverage of MNase fragment centres in +1 to +200bp window relative to TSS). C. Scheme of cofactor recruitment STAP-seq testing MED25 or GFZF Gal4 DNA-binding domain fusions recruited to a library of candidate promoter fragments. D. Box plot of the percent of STAP-seq signal (i.e. percent of initiation) originating at the dominant TSS at core promoters (CPs; N = 1266) and random regions (N = 639) that are activated to similar extent by both GFZF and MED25 recruitment. Cofactor recruitment STAP-seq data from (Haberle et al., 2019). *P≤0.01; Wilcoxon rank sum test. E. Histogram representing the distribution of the width of the initiation region (i.e. part of the oligo covered by STAP-seq signal) for CPs (N = 1266) and random regions (N = 639) upon recruitment of either MED25 or GFZF. P-values: Wilcoxon rank sum test. F. Cofactor recruitment STAP-seq tracks of GFP, MED25 and GFZF recruitment for examples of a core promoter and a random region that are activated by both cofactors. Endogenous initiation pattern in S2 cells (CAGE) is shown at the bottom. G. Scheme of Pol II PIC recruitment to the two types of developmental promoters (TATA-box and non-TATA-box containing DPE and INR motifs), which occurs through direct engagement between the transcription machinery and developmental promoter sequence motifs, resulting in narrow initiation patterns, whereas housekeeping promoters recruit Pol II through housekeeping DNA-binding proteins and intermediary cofactors that interact with TFIIA and Trf2, resulting in dispersed initiation.

In contrast, DRE- and Ohler 1-containing housekeeping promoters showed a dispersed distribution of CAGE signal in relation to DRE and Ohler 1 motifs, even for promoters that contain only a single motif occurrence (Figure 6A and Supplementary Figure 6A and 6B). Therefore, even though these motifs directly bind the DREF and M1BP factors, which can in turn recruit TFIIA, they do not instruct TSS position. We propose that the lack of strict motif positioning and initiation site at these housekeeping promoters is a direct result of weaker and less defined DNA-PIC interactions.

As transcription initiation at housekeeping promoters was not aligned to a sequence feature, we considered whether the promoter-proximal chromatin structure, especially the nucleosome-depleted region (NDR) or the +1 nucleosome might constrain initiation patterns. Although the CAGE signal is not strongly aligned with the +1 nucleosome at developmental promoters, housekeeping promoters exhibit a defined broad distribution of CAGE signal in the NDR immediately upstream of a strongly positioned +1 nucleosome (Figure 6B and Supplementary Figure 6D). These data show that initiation at housekeeping promoters occurs in a rather broad NDR upstream of the +1 nucleosome and suggest that the chromatin structure might be involved in determining TSS positions as previously proposed (Field et al., 2008; Rach et al., 2011).

If the dispersed initiation at housekeeping promoters results from a different mechanism of Pol II PIC recruitment, then transcriptional activation from the housekeeping-type TFIIA recruitment factors GFZF, Putzig and Chromator described above should always lead to more dispersed TSS patterns, irrespective of the promoter sequence. To test this systematically, we recruited the developmental-type coactivator MED25 and the housekeeping-type coactivator GFZF to a library of candidate promoters and analyzed the transcription initiation patterns (data from Haberle et al., 2019; Figure 6C). Although the two coactivators preferentially activate distinct sets of promoters (Haberle et al., 2019), 1266 promoters and 1268 random control sequences were activated sufficiently strongly by both coactivators to compare the respective initiation patterns (>4 fold induction over GFP with FDR<0.05) (Supplementary Figure 6C and 6G).

To systematically assess the initiation patterns across these fragments, we calculated the proportion of initiation events at the dominant TSS compared to the sum of all initiation events across the entire promoter fragment. On average across all core promoter fragments, initiation was at the dominant TSS for 55% of events after MED25 recruitment but only 42% after GFZF recruitment (p=1.6×10^-28^; Wilcoxon rank-sum test, Fig. 6D) and this difference persisted when housekeeping and developmental promoter sequences were analyzed separately (Supplementary Figure 6E) and even for random non-promoter fragments, for which the corresponding proportions were 59 vs. 49% (p= 2.4×10^-22^; Fig. 6D).

Consistently, when we examined all substantially activated TSSs within the non-promoter fragments (Supplementary Figure 6G), we found a single TSS for 47% of the fragments upon MED25 recruitment, while only 7% had 5 or more TSSs. In contrast, GFZF recruitment led to a single TSS for only 34% of the fragments, while 17% had 5 or more TSSs (Supplementary Figure 6F).

Moreover, MED25-induced transcription initiated for most promoters (51%) within a narrow 20bp region, while GFZF-induced transcription generally initiated in a much broader region of 30 to 75bp (only 24% promoters initiated within 20bp; Figure 6E). The existence of distinct initiation patterns at the same DNA sequence after MED25 versus GFZP recruitment is illustrated by the promoter of the Mcm3 gene and an intronic sequence within the DIP-kappa gene that does not initiate transcription endogenously (Figure 6F). The activation of transcription in characteristically different initiation patterns were also observed for two additional developmental (p300 and Lpt) and two housekeeping cofactors (Putzig and Chromator), respectively (Supplementary Figure 6H).

Thus, cofactor recruitment under identical conditions in an identical sequence context led to initiation patterns that are characteristically different for developmental cofactors (e.g. MED25) and housekeeping cofactors (e.g. GFZF), suggesting coactivators impose distinct patterns due to their different mechanisms of recruiting TFIIA, and the Pol II PIC.

## Discussion

In contrast to a prevalent model that Pol II PIC assembly and transcription activation occur similarly at all promoters, we find that different core promoter types recruit and activate Pol II via distinct strategies that depend on different factors.

Developmental promoter DNA is sufficient to recruit and assemble a Pol II PIC from nuclear extract *in vitro,* by having high affinity to GTFs such as TBP. Found as part of a soluble Pol II holoenzyme in yeast, TBP in complex with TFIIA is tightly associated with chromatin in metazoans and important in directing Pol II PIC assembly on DNA and cofactor mediated transcription *in vitro* (Kimura et al., 1999; Koleske and Young, 1995; Lieberman et al., 1997).

Our data indicates that most TATA-less promoters are independent of TBP and utilize TRF2, or TBP and TRF2 in a redundant fashion. Transcription in the absence of TBP has been observed for particular promoters (Wieczorek et al., 1998; Kwan et al., 2021) and cell types (Martianov et al., 2002; Gazdag et al., 2016), potentially involving TBP paralogs such as TRF2 in flies. Even though TRF2 has been reported to be unable to bind DNA directly (Rabenstein et al., 1999; Baumann et al., 2018), it may be recruited indirectly to promoters, potentially through interactions with TFIIA and/or TFIID (Baumann et al., 2017). This is analogous to transcription initiation during oocyte growth when the mammalian TBP paralog TBPL2 cooperates with TFIIA to initiate transcription independently of TFIID (Yu et al., 2020). The promoters of snRNA genes also function independently of TBP yet depend on SNAPc. At these promoters, SNAPc seems to directly bind TFIIA and/or TFIIB via an interface shared with TBP (Dergai et al., 2018; Mittal et al. 1999).

The partial redundancy of TBP and TRF2, especially when one of the two is depleted reconciles our results with recent structural studies of PIC assembly at non-TATA-box promoters (Chen et al., 2021): as TBPL1 or other TBP paralogs had not been considered during complex assembly *in vitro*, TBP was included in the PIC, irrespective of the promoter type. This might have been possible given the flexibility of the PIC, including TFIID that has been reported as sufficiently flexible to accommodate either TBP or TRF2 at different classes of promoters (Louder et al., 2016).

Housekeeping promoters bind sequence-specific transcription factors such as DREF and M1BP, which in turn interact with cofactors such as GFZF, Chromator and Putzig that – directly or indirectly – recruit GTFs (e.g. TFIIA) and Pol II (Figure 7G) (Baumann et al., 2018). These differences in the assembly and stability of the DNA-protein interface and protein complexes might explain the distinct transcription initiation patterns at developmental and housekeeping promoters, which generally exhibit focused and dispersed initiation patterns, respectively. Indeed, forced recruitment of housekeeping activators such as GFZF to arbitrary DNA sequences is sufficient to induce broad transcription initiation patterns, consistent with the initiation patters observed at housekeeping promoters *in vivo* and alternative PIC recruitment. This directly links the transcription activating cofactors of developmental and housekeeping programs to the distinct initiation patterns observed for the respective promoters.

The alternative mechanisms converge on TFIIA that is essential for transcription initiation at all promoter types. A central role of TFIIA recruitment for transcription initiation is consistent with the direct interaction of the TBP paralog TBPL2 with TFIIA in oocyte transcription (Yu et al., 2020), the direct interaction of SNAPc with TFIIA and/or TFIIB (Dergai et al., 2018) and non-canonical Pol II transcription of transposon-rich and H3K9me3-marked piRNA source loci in Drosophila germ cells through the TFIIA paralog moonshiner and TRF2 (Andersen et al., 2017). Essentially for some or all promoter types might extend to other GTFs that we could not test here, including TFIIB that is required at most promoters in human HAP1 cells (Santana et al., 2022).

Some of the features of Drosophila housekeeping promoters, including the dispersed patterns of transcription initiation, are similarly observed for the majority of vertebrate CpG island promoters comprising roughly 70% of all promoters (Carninci et al., 2006; Danks et al., 2018; FANTOM Consortium and the RIKEN PMI and CLST (DGT) et al., 2014; Saxonov et al., 2006). The functional regulatory dichotomy of these promoters combined with the evidence of distinct PIC composition and initiation mechanisms here and in other recent studies (Baek et al., 2021; Haberle et al., 2019) suggest that we need to challenge the notion of a universal model of rigid and uniform PIC assembly. It will be exciting to see future functional, biochemical and structural studies revealing more diverse transcription initiation mechanisms at the different promoter types in our genomes.

## Author Contributions

AS and LS conceived and designed the experiments and wrote the manuscript, with help from VH. LS, KB and OH generated endogenously tagged AID cell lines and performed PRO-seq. LP and AV performed analysis of PRO-seq data. FN & LS performed TFIIA immunoprecipitation for mass spectrometry. LS performed DNA affinity purifications. KM & ER performed mass spectrometry. AV & VL analyzed mass spectrometry data. VH analyzed Cut’N’Run, ChIP-seq and COF-STAP-seq data.

## Acknowledgements

We thank Ursula Schoeberl (IMP) and Maja Gehre (IMBA) for advice and help establishing PRO-seq and CUT&RUN and Clemens Plaschka (IMP), Carrie Bernecky (IST Austria), Dylan Taatjes (University of Colorado) and all members of the Stark lab for feedback and help on this project and manuscript. Next generation sequencing was done at the Vienna Biocenter Core Facilities GmbH (VBCF) Next-Generation Sequencing Unit (http://vbcf.ac.at); mass spectrometry was done by the mass spectrometry unit at IMP/IMBA/GMI. We thank Life Science Editors for comments on the manuscript. Research in the Stark group has been supported by the European Research Council (ERC) under the European Union’s Horizon 2020 research and innovation programme (grant agreement no. 647320) and by the Austrian Science Fund (FWF, F4303-B09 and P33157). Basic research at the IMP is supported by Boehringer Ingelheim GmbH and the Austrian Research Promotion Agency (FFG). LS is supported by a DOC PhD Fellowship from the Austrian Academy of Sciences. For the purpose of Open Access, the author has applied a CC-BY-NC-ND 4.0 International license to this pre-print.

## Resource availability

### Lead contact

Further information and requests for resources and reagents should be directed to and will be fulfilled by the lead contact, Alexander Stark.

### Materials availability

All cell lines and plasmids generated in this study are available upon request.

### Data and code availability

PRO-seq and CUT&RUN data have been deposited to the Gene Expression Omnibus (GEO), accession GSE181257, which will be publicly available at the date of publication. Raw mass spectrometry data of DNA-affinity purification have been deposited to ProteomeXchange through the PRIDE server under identifier PXD028090, and mass spectrometry data of TFIIA-L immunoprecipitation under identifier PXD028094.

## Experimental model and subject details

### Cell lines

Drosophila Schneider S2 cells were purchased from Gibco and were maintained according to the manufacturer’s recommendations as described in the method details. Engineered cell lines were created using CRISPR/Cas9 as described in the method details.

## Method details

### Cell culture

Drosophila melanogaster S2 cells were obtained from Thermo Fisher and maintained in Schneider’s Drosophila Medium supplemented with 10% heat-inactivated fetal bovine serum.

### Generation of endogenously tagged AID cell lines

A parental cell-line expressing the osTir ligase was created with a knock-in approach by introducing a vector expressing a gRNA/Cas9 targeting the carboxyl terminus of the Act5C, with a P2A before the osTir-mCherry construct, leading to constitutive expression of the osTir ligase. Wild type S2 cells were electroporated using the MaxCyte-STX system at a density of 1×10^7^ cells per 100µl and 20 µg of DNA using the pre-set protocols. Cells were selected with puromycin and FACS sorted based on mCherry fluorescence into individual 96-well plates to generate individual clones which were screen by PCR and for their ability to degrade transfected AID-tagged proteins. To generate AID cell lines we have electroporated a knock-in cassette to either the N-terminal or C-terminal of the gene of interest, a cassette containing a mAID-3xFLAG tag. Cells were electroporated as described above. Electroporated cells were selected on 5 µg/mL blasticidin and diluted to individual 96-well plates to generate single clones. Single clones were amplified and genotyped using a PCR to the presence of a homozygous knock-in and confirmed with Sanger sequencing. To generate a double tagged TBP + TRF2 AID cell line, the TRF2 AID cell line was electroporated with a knock-in cassette containing a TBP-AID with a hygromycin selection marker. Cells were selected for one week on 5 µg/mL hygromycin and single clones were generated as above. Single clones were additionally tested for their ability to degrade the AID-3xFLAG tagged proteins on a western blot using an anti-FLAG antibody.

### Selection of promoters and controls for DNA-affinity purification

Unique CAGE corrected TSSs were scored for PWM for core-promoter motifs (TATA-box, INR, DPE, TCT, Ohler1/6, DRE), and the highest scoring TSSs that were expressed in Drosophila S2 cells (≥5tpm), and were inducible in STAP-seq (Arnold et al., 2016) were used. The following PWM score cut-offs were used for the selected motifs: TCT ≥ 95%, Ohler 1+6 ≥ 95%, DRE 100%, TATA-box and INR ≥ 95%, DPE and INR ≥ 95%, and INR only ≥ 95%. Length matched control regions were selected from the Drosophila genome to not overlap transcribed regions, and did not show any sign of transcription in any Drosophila developmental CAGE data. Selected promoters are listed in supplementary table 1.

### Cloning promoter constructs

Promoter regions were PCR amplified from S2 cell genomic DNA using primers containing gibson overhangs corresponding to the BglII and HindII restriction sites on pGL3 with Q5 high-fidelity 2x master mix (NEB). PCR products were cleaned with AMPURE beads and eluted in water. Gibson reactions were performed with a Gibson assembly master mix (NEB) according to the manufacturer’s recommendations. 1ul of Gibson reaction was electroporated into MegaX DH10B electrocompetent cells (Thermo). Single clones were picked and grown in 5mL bacterial cultures. Minipreps were performed using a Qiagen kit, and Sanger-sequencing was performed in-house. Correct plasmid clones were used as a template for amplification of biotinylated DNA.

### Preparation and immobilization of biotinylated DNA

Biotinylated DNA was generating using a forward primer containing a Biotin TEG group on the 5’ end obtained from Sigma Aldrich: Biotin TEG 5’, and a reverse Reverse 3’ primer (see resource table for primer sequences). At least 2mL of total PCR volume (performed in 50ul reactions) for each individual promoter sequence was amplified individually for each replicate. PCR reactions were pooled and DNA was purified using AMPURE beads and eluted in water. For each sample, 50µl of Dyna M280 Streptavidin were used and coupled to 15µg of cleaned biotinylated PCR product according to the manufacturer’s recommendations. The beads were placed in an equivalent volume of DBB (150 mM NaCl, 50 mM Tris/HCl pH, 8.0, 10 mM MgCl2) and used immediately for DNA-affinity purification assay.

### Preparation of nuclear extracts

Nuclear extracts from drosophila S2 cells were prepared as previously described with the following modifications (Dignam et al., 1983). Three billion drosophila S2 cells were harvested by resuspension and washed with PBS. The cell pellet was resuspended in buffer A (10mM HEPES pH7.9, 1.5mM MgCl2, 10mM KCl, 0.5mM DTT added fresh before use, and oComplete EDTA-free protease inhibitors) placed on ice for 10 minutes. Cells were spun down at 700g for 5 minutes, supernatant removed, and cells were resuspended in 5 cell-pellet volumes of buffer A supplemented with 0.5% NP-40. Cell suspension was dounced in a Beckman 15mL dounce with a ‘loose’ pestle for 10 strokes to isolate nucli. Cells were spun down at 2000g for 5 minutes at 4°C, supernatant containing the cytoplasmic fraction was removed, and cell pellet containing the nuclei was resuspended in three pellet volume of buffer C (0.5M NaCl, 20mM HEPES pH7.9, 25% glycerol, 1.5mM MgCl2, 0.2mM EDTA, 0.5mM DTT added before use, oComplete EDTA-free protease inhibitors), and placed over a 10% sucrose cushion made in buffer C, and spun down at 3000g for 5 minutes at 4°C. Supernatant was removed and the pellet was resuspended in buffer C, equivalent of 1mL per 1 billion starting cells. Nuclei were dounced in a Beckman 7mL dounce with a “tight” pestle for 20 strokes. Lysed nuclei were rotated at 4°C for 30 minutes, and then spun down at 20,000g for 10 minutes at 4°C. The supernatant was the soluble nuclear fraction was dialyzed in buffer D (20mM HEPES pH7.9, 20% glycerol, 0.1M KCl, 0.2mM EDTA, 0.5mM DTT added before use, and oComplete EDTA-free protease inhibitors) using Slide-A-Lyzer dialysis casettes with a 3.5kD molecule weight cut-off for 6 hours with two buffer exchanges. Protein concentration of the nuclear extract was determined with a Qubit protein assay kit according to the manufacturer’s instructions. Dialyzed nuclear extract was snap frozen in liquid nitrogen and stored at −80°C until use.

### DNA-affinity purification and on-bead digest

50ul of DNA-immobilized beads were mixed with 400µg of nuclear extract and 1200ng sheared salmon sperm DNA in Axygen 1.5mL tubes. Reactions were incubated at room temperature for 40 minutes with rotation. Beads were then magnetically pelleted, washed once with buffer DBB (supplemented with 0.5%NP-40), and resuspended in DBB supplemented 0.75% formaldehyde for 10 minutes at room temperature with rotation. Beads were resuspended in 50 µl of 100mM ammonium bicarbonate. 600ng of Lys-C (Wako) was added to the beads and digests were incubated at 37°C for 4 hours in a thermoblock with shaking at 800rpm. Beads were magnetically pelleted, and the supernatant was transferred to a new 0.6mL Axygen tube. Samples were incubated with 6µl of a 6.25mM TCEP-HCl solution (Sigma) at 60°C for 30 minutes in a thermoblock with rotation at 400rpm. Next, 6µl of 40mM MMTS was added and incubated for 30 minutes in the dark. Finally, 600ng of trypsin gold (Promega) was added and digests were incubated at 37°C overnight. Digests were stopped with 10µl of 10% TFA solution. 30% of the reaction volume was used for Nano LC-MS/MS analysis. Results from the promoter DNA-affinity purification mass spectrometry are listed in supplementary table 1.

### Nano LC-MS/MS Analysis for DNA-affinity purification

An UltiMate 3000 RSLC nano HPLC system (Thermo Fisher Scientific) coupled to a Q Exactive HF-X equipped with an Easy-Spray ion source (Thermo Fisher Scientific) or an Exploris 480 mass spectrometer equipped with a Nanospray Flex ion source (Thermo Fisher Scientific) was used. Peptides were loaded onto a trap column (PepMap Acclaim C18, 5 mm × 300 μm ID, 5 μm particles, 100 Å pore size, Thermo Fisher Scientific) at a flow rate of 25 μl/min using 0.1% TFA as mobile phase. After 10 min, the trap column was switched in line with the analytical column (PepMap Acclaim C18, 500 mm × 75 μm ID, 2 μm, 100 Å, Thermo Fisher Scientific). Peptides were eluted using a flow rate of 230 nl/min, and a binary linear 3h gradient, respectively 225 min.

The gradient started with the mobile phases 98% A (0.1% formic acid in water) and 2% B (80% acetonitrile, 0.1% formic acid), increased to 35% B over the next 180 min, followed by a steep gradient to 90%B in 5 min, stayed there for 5 min and ramped down in 2 min to the starting conditions of 98% A and 2% B for equilibration at 30°C.

### TFIIA immunoprecipitation

Drosophila S2 cells endogenously tagged with an AID-3xFLAG were used for the bait, while the parental background cells only expression the osTir ligase were used as a control immunoprecipitation. Lysates were generated from 500 million cells. Cells were washed in PBS and pelleted by centrifugation. Cell pellet was resuspended in 10mL of hypotonic swelling buffer (10mM Tris pH7.5, 2mM MgCl2, 3mM CaCl2, protease inhibitors) and incubated for 15 minutes at 4°C. Cells were centrifuged for 10 minutes at 700g and at 4°C. Cells were resuspended in 10mL of GRO lysis buffer (10mM Tris pH7.5, 2mM MgCl2, 3mM CaCl2, 0.5% NP-40, 10% glycerol, 1mM DTT, protease inhibitors) and rotated for 30 minutes at 4°C. Nuclei were centrifuged at 700g and at 4°C. Supernatant was removed and nuclei were resuspended in 1mL of IP lysis buffer (100mM NaCl, 20mM HEPES pH7.6, 2mM MgCl2, 0.25% NP-40, 0.3% Tirton X-100, 10% glycerol) and rotated for 30 minutes at 4°C. Lysed nuclei were centrifuged for 5 minutes at 20000g at 4°C. The supernatant containing the soluble nucleoplasm was kept. While the chromatin pellet was resuspended in a 300mM NaCl IP lysis buffer (300mM NaCl, 20mM HEPES pH7.6, 2mM MgCl2, 0.25% NP-40, 0.3% Tirton X-100, 10% m glycerol) and sonicated Diagenode Bioruptor sonicator: 10 min (30 sec on/30 sec off) at low intensity. The sheared chromatin was centrifuged as before and the soluble supernatant was removed and mixed with the soluble nucleoplasmic fraction. The resulting mixture was centrifuged again for 5 minutes at 20000g at 4°C to remove insoluble proteins. Anti-FLAG M2 beads (Sigma-Aldrich) were equilibrated by three 10 minute washes with 150mM NaCl IP lysis buffer, and resuspended back in their original volume. Immunoprecipitation reactions were set up with 50ul of Anti-FLAG beads and 1mg of the nuclear lysates overnight with rotation at 4°C. Immunoprecipitation reactions were magnetically pelleted and washed with 150mM IP lysis buffer three times, 10 minutes each with rotation at 4°C. Next, to remove detergent, the reactions were washed 4 times, 10 minutes each at 4°C with a no-detergent buffer (130mM NaCl, 20mM Tris pH7.5). Reactions were resuspended in 50µl of 100mM ammonium bicarbonate and on-bead tryptic digest was carried out as described in the DNA-affinity purification and on-bead digest section. Results of the TFIIA-L immunoprecipitation are listed in supplementary table 2.

### Nano LC-MS/MS Analysis for TFIIA-L Immunoprecipitation

A Q Exactive HF-X mass spectrometer was operated in data-dependent mode, using a full scan (m/z range 380-1500, nominal resolution of 60,000, target value 1E6) followed by MS/MS scans of the 10 most abundant ions. MS/MS spectra were acquired using normalized collision energy of 27, isolation width of 1.4 m/z, resolution of 30.000, target value of 1E5, maximum fill time 105ms. Precursor ions selected for fragmentation (include charge states 2-6) were put on a dynamic exclusion list for 60 s. Additionally, the minimum AGC target was set to 5E3 and intensity threshold was calculated to be 4.8E4. The peptide match feature was set to preferred and the exclude isotopes feature was enabled.

### LC-MS/MS analysis for TFIIA-L immunoprecipitation

The Orbitrap Exploris 480 mass spectrometer (Thermo Fisher Scientific), was operated in data-dependent mode, performing a full scan (m/z range 380-1200, resolution 60,000, target value 3E6) at 2 different CVs (−50, −70), followed each by MS/MS scans of the 10 most abundant ions. MS/MS spectra were acquired using a collision energy of 30, isolation width of 1.0 m/z, resolution of 45.000, the target value of 1E5 and intensity threshold of 2E4 and fixed first mass of m/z=120. Precursor ions selected for fragmentation (include charge state 2-5) were excluded for 30 s. The peptide match feature was set to preferred and the exclude isotopes feature was enabled.

### Mass-Spectrometry Data Processing

For peptide identification, the RAW-files were loaded into Proteome Discoverer (version 2.5.0.400, Thermo Fisher Scientific). All hereby created MS/MS spectra were searched using MSAmanda v2.0.0.16129 (Dorfer V. et al., J. Proteome Res. 2014 Aug 1;13(8):3679-84). RAW-files were searched in 2 steps: First, against the drosophila database called dmel-all-translation-r6.34.fasta (Flybase.org, 22,226 sequences; 20,310,919 residues), or against an earlier version dmel-all-translation-r6.17.fasta (21,994 sequences; 20,118,942 residues) / a small custom drosophila database (107 sequences; 61,976 residues), each case supplemented with common contaminants, using the following search parameters: The peptide mass tolerance was set to ±5 ppm and the fragment mass tolerance to ±15 ppm (HF-X) or to ±6 ppm (Exploris). The maximal number of missed cleavages was set to 2, using tryptic specificity with no proline restriction. Beta-methylthiolation on cysteine was set as a fixed modification, oxidation on methionine was set as a variable modification, the minimum peptide length was set to 7 amino acids. The result was filtered to 1 % FDR on protein level and was used to generate a smaller sub-database for further processing. As a second step, the RAW-files were searched against the created sub-database using the same settings as above plus the following search parameters: Deamidation on asparagine and glutamine were set as variable modifications. In some data sets acetylation on lysine, phosphorylation on serine, threonine and tyrosine, methylation on lysine and arginine, di-methylation on lysine and arginine, tri-methylation on lysine, ubiquitinylation residue on lysine, biotinylation on lysine, formylation on lysine were set as additional variable modifications. The localization of the post-translational modification sites within the peptides was performed with the tool ptmRS, based on the tool phosphoRS (Taus et al., 2011). Peptide areas were quantified using the in-house-developed tool apQuant (Doblmann et al., 2018). Proteins were quantified by summing unique and razor peptides. Protein-abundances-normalization was done using sum normalization. Statistical significance of differentially expressed proteins was determined using limma (Smyth, 2004).

### PRO-seq

PRO-seq was performed according to (Mahat et al., 2016) with the following modifications. 10 million Drosophila Schneider S2 cells were used for each replicate, spiked in with 1% human HCT116 cells. Cells were harvested by centrifugation and cells were permeabilized with cell permeabilization buffer (10mM tris Ph 7.5, 300mM sucrose, 10mM CaCl2, 5mM MgCl2, 1mM EGTA, 0.05% tween-20, 0.1% NP-40, 0.5mM DTT, supplemented with protease inhibitors). Permeabilization was carried by resuspending the cells in 10mM of permeabilization buffer and spinning down the cells for a total of three buffer exchanges. Nuclei were resuspended in 100µl of storage buffer (10mM tris pH 7.5, 25% glycerol, 5mM MgCl2,0.1mM EDTA and 5mM DTT) and snap frozen in liquid nitrogen for later use, or immediately proceeded to the run-on reaction. Nuclear transcription run-on was carried by adding 100µl of a 2x run-on buffer (10mM tris pH8, 5mM MgC2, 1mM DTT, 300mM KCl, 0.25mM ATP, 0.25mM GTP, 0.05mM Biotin-11-CTP, 0.05mM Biotin-11-UTP, 0.8U/µl murine RNase inhibitor, 1% sarkosyl) and incubated at 30C for 3 minutes. Reaction was terminated by adding 500ul Trizol-LS. Extraction was performed by adding 130µl of chloroform, after vortexing and centrifugation the aquesous fraction was kept and precipitated with 2.5 volumes of 100% ethanol and 1µl of glycoblue. The pellet was washed with 80% ethanol, air dried and resuspended in 50µl of water. RNA was denatured at 65C for 40 seconds before base hydrolysis with 5µl 1N NaOH for 15 minutes. Hydrolysis was quenched with 25 µl of 1M tris-HCl pH6.8. Samples were purified on a Bio-Rad P30 column. Biotinylated nascent RNA was recovered by incubating with 50µl of M280 streptavidin beads for 30 minutes at room temperature with rotation. Beads were washed twice each with high salt buffer (2M NaCl, 50mM Tris pH 7.5, 0.5% Tirton X-100) and binding buffer (300mM NaCl, 10mM Tris pH 7.5, 0.1% Tirton X-100) and once with low salt buffer (5mM Tris pH 7.5, 0.1% Tirton X-100). RNA was extracted off the beads using Trizol and cleaned on a Direct-zol column (Zymo). RNA was eluted from the column using 5 µl the 3’ RNA linker. Overnight ligation at 16°C was performed with T4 RNA ligase I. The following day biotinylated RNA was recovered with 50µl of M280 streptavidin beads for 30 minutes at room temperature and washed as described. The RNA was treated with CapCLIP Pyrophosphatase (Biozyme) on the beads for 1 hour at 37°C, followed by T4 polynucleotide kinase (NEB) for 1 hour at 37°C. Beads were washed as described and an on-bead ligation was set up with T4 RNA ligase I and the 5’ RNA linker at room temperature with rotation at 4 hours. Next, the beads were washed as described and the RNA was extracted off the beads with 300µl Trizol and purified on a Direct-zol column, eluted in water. Eluted RNA was used for reverse transcription with Superscript III Reverse Transcriptase (Thermo) according to the manufacturer’s recommendations. Half of the reverse transcription reaction was used for amplification with a KAP real-time PCR mixture (KAPA Biosystems) using the Illumina Truseq small RNA library amplification kit primers. Libraries were amplified in 8-12 cycles. Primer dimers were removed from the libraries with AMPURE beads and sent for next-generation sequencing.

### PRO-seq data mapping

PRO-seq libraries were sequenced to a depth of 3.8 - 38.9 million reads using single-end sequencing and read length of 50 bp. We used unique molecular identifiers (UMIs) to distinguish between PCR duplicated identical reads and reads stemming from distinct RNA molecules with an identical sequence. The latter will have identical sequences but different UMIs and therefore allows more accurate quantification of transcripts. RNA oligos containing UMIs of 8-10 nt in length were ligated to the 3’ end of all reads before PCR amplification and then computationally removed to prevent interference during genome alignment. Cutadapt 1.18 (Martin et al., 2011) with default options was used to find and trim the sequencing adapter at the 3’ end and filtered for reads ≥ 10 nts long. Only after read alignment we corrected for PCR duplicated transcripts and to more accurately quantified transcripts: reads containing the same sequence and reads aligning to the same genomic position were collapsed to unique UMIs.

To align reads, we generated an artificial genome containing sequences for tRNAs and rRNAs only, which allows for noise reduction of short reads aligning to multiple positions. Next, all unmapped reads were captured using samtools version 1.9 (Li et al., 2009) with -f 4 option, which were then aligned to the *D. melanogaster* reference genome BDGP R5/dm3. Following this, reads not aligning to the dm3 genome were aligned to the *H. sapiens* reference genome GRCh37/hg19 (used as spike-in). For genome alignment we used bowtie version 1.2.2 (Langmead et al., 2009) allowing two mismatches (-v 2). For alignment to the artificial genome we allowed reads having up to 1000 reportable alignments, but reporting only the best alignment (-m 1000 --best --strata) to meet the highly repetitive and conserved nature of tRNAs and rRNAs. Alignment to the reference genomes was run allowing only reads aligning uniquely (-m 1).

We generated an artificial genome containing the ribosomal RNA primary transcript CR45847 (http://flybase.org/reports/FBgn0267507), all annotated tRNA genes from Dmel 5.57 and tRNAs predicted from Genomic tRNA database, published 2009, http://lowelab.ucsc.edu/GtRNAdb/ (accessed August 17th, 2020; http://lowelab.ucsc.edu/download/tRNAs/eukaryotic-tRNAs.fa.gz). We used R packages GenomicRanges 1.34.0 (Lawrence et al., 2013), Biostrings 2.50.2 (https://bioconductor.org/packages/Biostrings) and BSgenome.Dmelanogaster.UCSC.dm3 1.4.0 (Team TBD (2014). *BSgenome.Hsapiens.UCSC.hg17: Full genome sequences for Homo sapiens (UCSC version hg17)*. R package version 1.3.1000.

Since application of the usual PRO-seq protocol delivers reads corresponding to the reversed complement of the nascent RNA, the reads aligning to the minus strand originated from transcripts with the sequence on the plus strand and vice versa. Additionally, only the end of the transcript where RNA Pol II was actively transcribing was included for the downstream analysis. Reads were switched and shortened accordingly using the bigBedtoBed utility (Kent et al., 2010).

### ChIP-seq and ChIP-exo data analysis

ChIP-seq and ChIP-exo data sets were taken from (Baumann and Gilmour, 2017; Gurudatta et al., 2013; Shao and Zeitlinger, 2017). Coverage was calculated over a 1-kb window centered on the TSS of each promoter type. Data was normalized for the transcription level as measured by PRO-seq, which was further normalized by gene length for each individual promoter.

### Promoter motif annotations

We generated an R table in version 3.5.3 (R Core Team, 2019) containing transcripts of all protein-coding genes and corrected their transcription starting site with hits supported by CAGE experiments (Brown et al., 2014). First, TSSs were corrected by CAGE signal from S2 cells downloaded from modENCODE 5331 that lie within a window of ±250 bps. If no hit was found, CAGE signals from mixed embryos or a developmental timecourse from modENCODE 5338-5348, 5350 and 5351 were used within the same window. If the TSS was left unsupported we repeated this using a ±500 bp window or kept the annotated TSS. We kept the longest transcript per unique TSS. We used the R packages CAGEr 1.24.0 (Haberle et al., 2014) and GenomicRanges 1.34.0(Lawrence et al., 2013). Additionally, all transcripts were annotated with known *D. melanogaster* core promoter motifs as described in a previous study (Haberle et al., 2019) with small changes regarding match thresholds for TATAbox to 90%, DPE to 98% DRE to 98% and Ohler6 to 97%. TCT motif was further limited to ribosomal proteins.

### Generation of browser tracks of PRO-seq data

For visualization of PRO-seq data we converted bigBed files to bigWig files using kentUtils bigBedToBed utility (Kent et al., 2010), normalized by the number of reads aligned to dm3 (and considered number of reads aligned to hg19 for TFIIA samples) and calculated the coverage using genomeCoverageBed from bedtools 2.27.1 (Quinlan and Hall, 2010) before converting to a bigWig file using KentUtils wigToigWig utility. BigWig files were visualized with the UCSC Genome Browser (Kent et al., 2010).

### Differential expression

Differential expression was calculated using the DESeq function from the DESeq2 package v.1.30.1 (Love et al., 2014) providing the normalization factors as sizeFactors. Normalization factors were calculated based on the number of reads aligned to *D. melanogaster* reference genome and for TFIIA quantified spike-in reads were additionally considered. We used Benjamini-Hochberg adjusted p-values to determine significantly deregulated transcripts.

### CUT&RUN

CUT&RUN was performed as previously described (). Briefly, half a million cells per replicate were washed with PBS, and resuspended in 1.5 mL Wash Buffer (20mM HEPES pH 7.3, 150mM NaCl, 0.5mM Spermidine, Roche complete Inhibitor, EDTA-free). Cells were washed three times total with Wash buffer, and resuspended in 10uL of ConcavadinA magnetic beads equilibrated in Binding buffer (20mM HEPES pH 7.3, 10mM KCl, 1mM CaCl2, 1mM MnCl2, Roche complete Inhibitor, EDTA-free). Cells were mixed with the beads for 10 minutes following resuspension in 50uL Antibody buffer (20mM HEPES, 150mM NaCl, 0.5mM Spermidine, 0.025% Digitonin, 2mM EDTA, Roche Complete Inhibitor). M2 FLAG antibody was added to a final dilution of 1:100, and incubated overnight at 4°C. Beads were washed 3x and resuspended in 150uL Digitonin Buffer (20mM HEPES, 150mM NaCl, 0.5mM Spermidine, 0.025% Digitonin, Roche Complete Inhibitor). pAG-MNase was added to a final concentration of 700 ng/mL and incubated with rotation for one hour at 4°C. Cells were washed 3 times with Digitonin Buffer supplemented with calcium chloride to final concentration of 2mM, and rotated for 2 hours at 4°C. Digestion was stopped by the addition of 100uL STOP buffer (340 mM NaCl, 20mM EDTA, 4mM EGTA, 0.02% Digitonin, 50ug/mL RNase A, 50 ug/mL Glycogen). Samples were incubated for 10 minutes at 37°C with shaking at 500rpm on a thermocycler tabletop heatblock, followed by centrifugation for 5 minutes at 16000g. The supernatant was taken and digested with Proteinase K for 2 hours at 55°C and DNA extracted using a Qiagen PCR purification kit. The resulting DNA was cloned into a library using the NEBNext Ultra II DNA Library Prep Kit for Illumina libraries.

### STAP-seq data analysis of initiation events

Cofactor recruitment STAP-seq data from (Haberle et al., 2019) was analyzed at single nucleotide resolution counting unique transcripts initiated at each position in each tested oligo. The dominants TSS was determined as the position with the highest count, and the relative count was calculated by dividing the count at the dominant TSS with the total count for each oligo. To determine the number of activated TSSs in each oligo, the count at each position was divided by the count at the dominant TSS, and only the positions with a ratio of more than 20% were counted as activated TSSs.

### Defining Drosophila melanogaster promoter types

A set of ∼17,000 promoters of protein coding genes was classified into 9 groups based on PWM scores for the different CP motifs. The data was clustered with k-means to get the representative groups defined by the occurrence of specific motifs or motif combinations.

### Aligning CAGE data to promoter motif positions and +1 nucleosome centers

For the above defined promoter groups the positions of the defining CP motifs were determined relative to the dominant CAGE TSS (if they occurred within +/- 120 bp). Only promoters with a single occurrence of each motif were considered, and the position of the motif was used as a reference point to generate average plots of CAGE data. MNase-seq data from(Chereji et al., 2016), CAGE data from mixed embryos (Hoskins et al., 2011)

MNase-seq data was used to determine the position of the +1 nucleosome by taking the centers of MNase fragments between 100 and 200 bp long, calculating the coverage of such centers, and determining the position with the highest coverage in the region 150 bp downstream of the dominant CAGE TSS. These +1 nucleosome centers were used as a reference to generate average plots of CAGE data for each promoter group.

### Luciferase assay

Drosophila Schneider S2 cells were plated in 96-well plates, 1×10^5^ cells per well. Cells were transfected with 100ng of luciferase plasmid contaiing a DRE promoter or negative control sequence upstream of the luciferase gene, and 100ng of a plasmid containing renilla luciferase as a transfection efficiency normalization control using Lipofectamine 2000. Cells were lysed 48 hours after transfection with 50µl passive lysis buffer for 30 minutes at room temperature with shaking. Lysates were further diluted 10 fold in passive lysis buffer. 10µl of the diluted lysate was placed in 96-well plates compatible with luminescence read-out and measured with the Promega dual-luciferase assay kit according to the manufacturer’s recommendation on a BioTek Synergy H1 plate reader.

For COF recruitment luciferase assay in AID cell lines, we have first transfected the luciferase reporter and Gal4-COF expressing plasmids. After 24 hours we added 500uM auxin and waited an additional 24 hours prior to measurement of the luciferase signal.

## Supplementary Figure Legends

**Figure S1.**
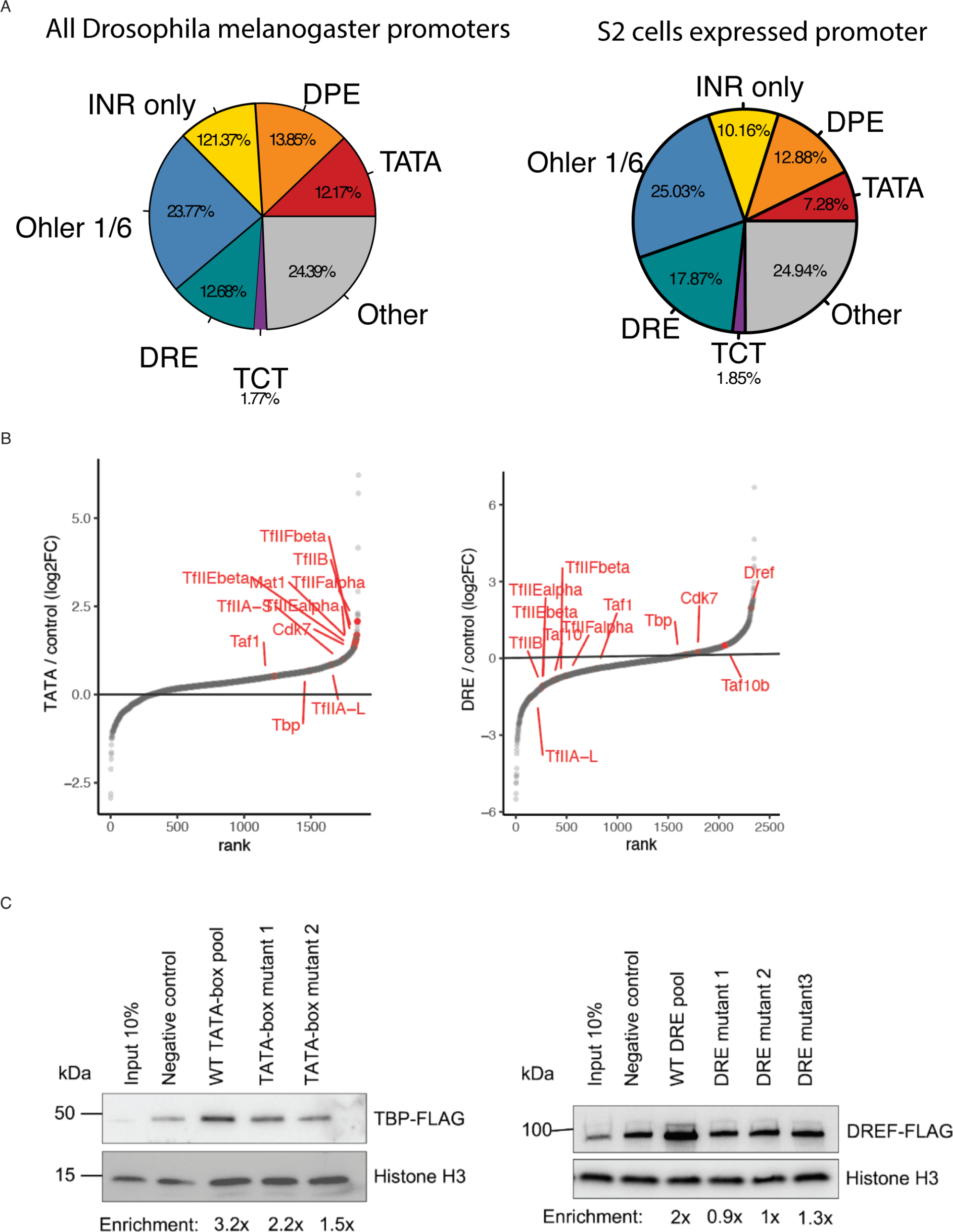
DNA affinity purifications uncover differentially bound proteins at functionally distinct promoters. A. Pie chart of all expressed Drosophila melanogaster protein coding gene promoters (∼170000) grouped based on motif content (left), and all expressed protein coding genes from Drosophila S2 cells (∼10000). Only the main motif groups studied in this paper that are classified as housekeeping or developmental are shown. Group labeled as “other” contains promoters with motifs such as Ohler 8 and E-box or not motifs which could not be assigned as developmental or housekeeping. B. Rank plot of protein binding enrichment on TATA and DRE promoters over the control DNA pool from the DNA-purification mass spectrometry assay. Highlighted proteins are the Pol II PIC components and the DRE binding factor DREF. C. DNA-purification assay with a pool of 25 TATA-box promoters, and two individual TATA-box promoters in which the TATA-box was mutated (left panel). The assay was performed with a nuclear extract expressing TBP-FLAG that was tracked with a western blot. DNA-purification of a pool of 20 DRE promoters and three individual DRE promoters in which the DRE motif was mutated. The assay was performed with a nuclear extact expressing DREF-FLAG and followed with a western blot (right panel).

**Figure S2.**
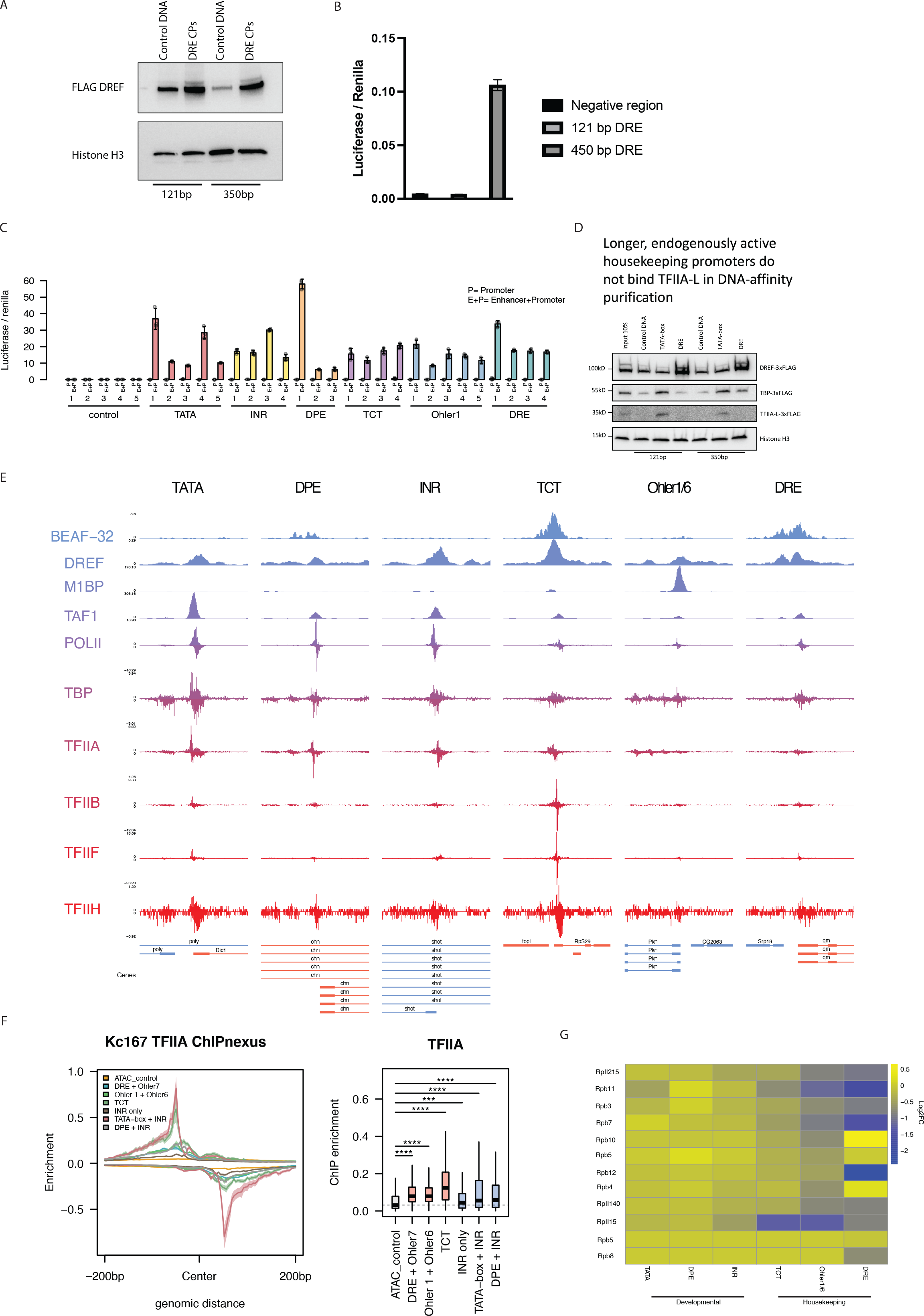
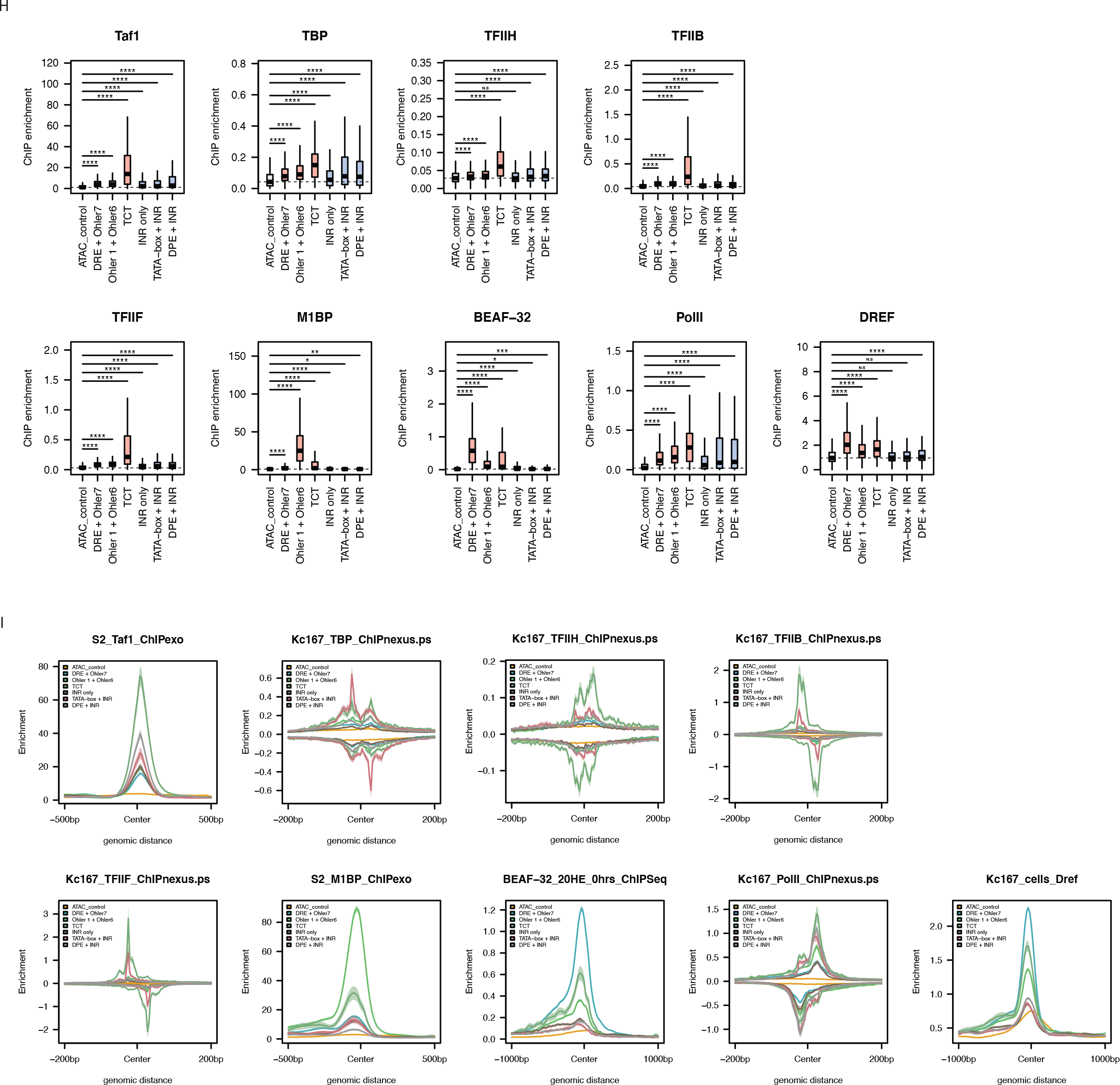
Developmental and housekeeping promoters bind different sets of proteins and GTFs. A. Elution fractions from the DNA-purification assay with a pool of 20 121bp or a pool of 10 350bp DRE promoters and length matched negative controls were performed with a nuclear extract expressing DREF-AID-3xFLAG tag and blotted for an anti-FLAG antibody. Both promoter lengths are able to enrich for DREF binding. B. Luciferase assay performed with DRE promoter fragments from the zip gene promoter that are 121bp or 450bp in length and a negative control region which is 450bp in length. Firefly luciferase values were normalized to co-transfected renilla luciferase values. C. Luciferase activity assay of 121bp long promoter fragments cloned upstream of a luciferase gene (P), or cloned in addition upstream of the Drosophila Zdfh1 enhancer (E+P). Firefly luciferase values were normalized to co-transfected renilla luciferase values. D. DNA-purification assay with pool of 121bp or 350bp long promoters with a nuclear extract expressing DREF-FLAG, TBP-FLAG and TFIIA-L-FLAG was followed by western blot with an anti-FLAG antibody. This direct comparison indicates that both short (inactive) and longer (endogenously active) housekeeping promoters are able to bind and enrich for sequence specific TFs such as DREF, however cannot bind TFIIA-L *in vitro*. E. Representative browser tracks of published ChIP-seq data of GTFs and promoter binding TFs (M1BP, DREF, BEAF-32) on the 6 different tested promoter types in this study. F. Meta-plot of TFIIA-L ChIP-seq data from panel E at the 6 different tested promoter types indicating TFIIA binds all active promoter types, although less strongly to housekeeping promoters and in a more dispersed fashion relative to the TSS (center). Box plot quantification of TFIIA ChIP-seq data at -/+ 200bp around the TSS. G. Heat map of log2FC values of DNA-affinity purification values for RNA polymerase II subunits across the six different promoter types tested. H. Box plots representing ChIP-seq signal of available GTFs and sequence specific TFs on the six different promoter types tested from Drosophila melanogaster embryos centered on the TSS (−200bp to +200bp). *Wilcoxon p<0.05. I. Meta plot of ChIP-seq signal of available GTFs and sequence specific TFs (as in panel H) on the six different promoter types centered on the TSS.

**Figure S3.**
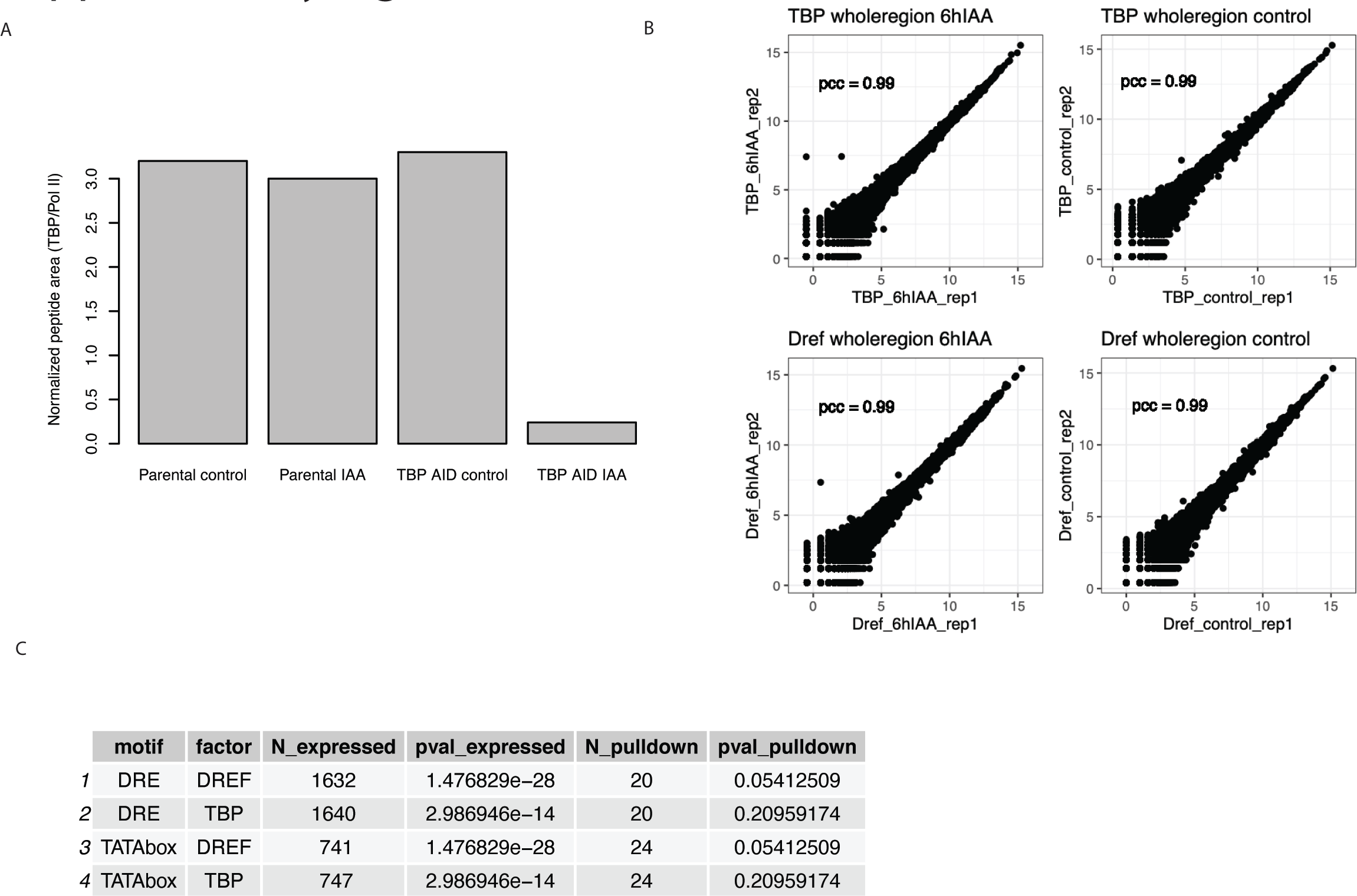
TBP and DREF are required by distinct sets of promoters. A. Mass-spectrometric quantification of TBP protein abundance in the parental cell line expressing the Tir1 ligase and the TBP N-terminally tagged AID cell line after 6 hours of 500uM auxin treatment. Normalize peptide abundance from label free mass-spectrometric quantification indicates roughly 3% of TBP remains after auxin treatment as compared to the control. B. Pearson correlation of PRO-seq signal along the promoter and gene body region of all protein-coding transcripts using library-normalized reads between biological replicates. Correlation coefficient displayed. C. The number of DRE and TATA-Box expressed promoters in each of the DREF and TBP AID tagged cell lines. P-value calculated with FDR indicate down-regulation of TATA-Box or DRE promoters compared with all expressed promoters.

**Figure S4.**
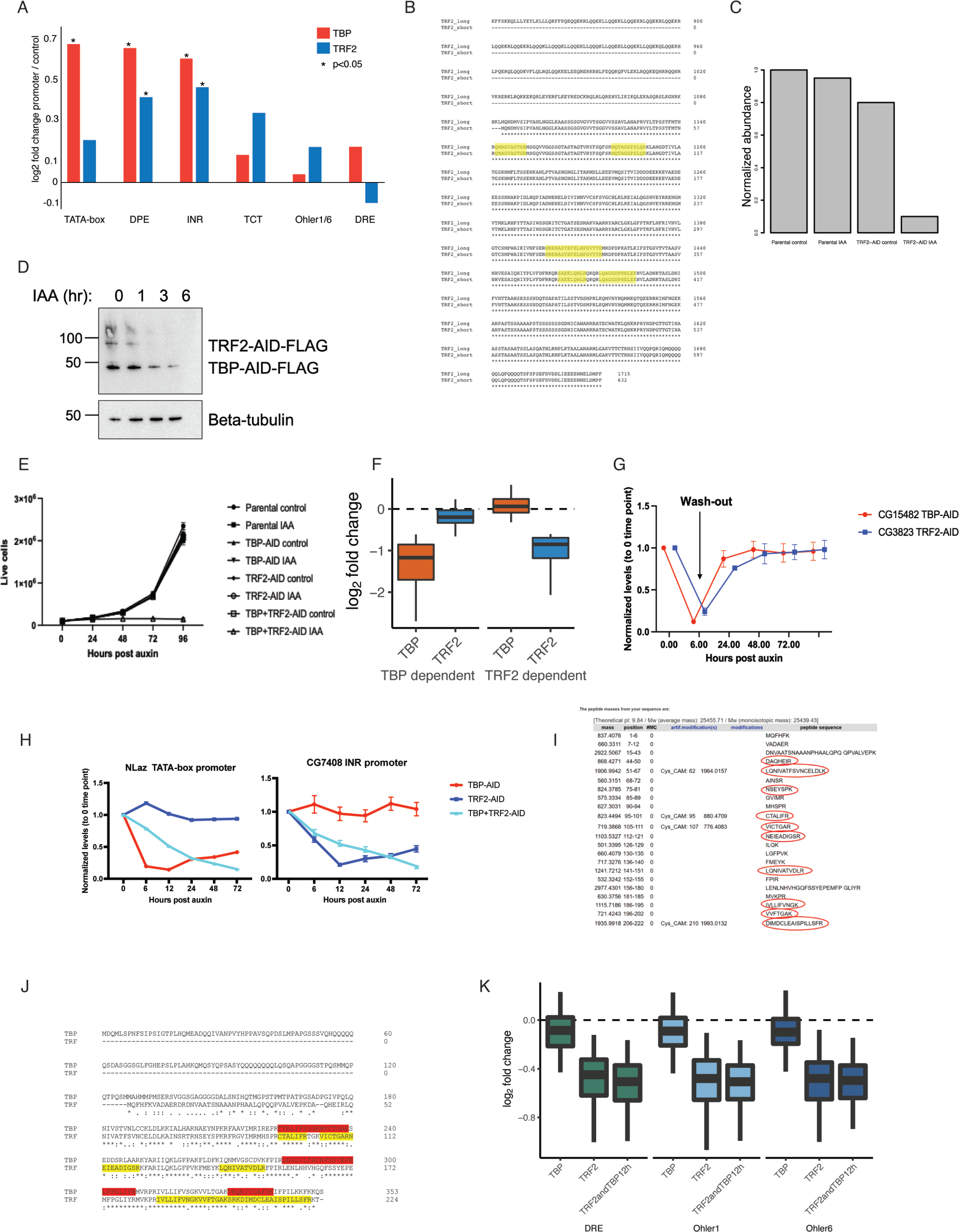
TBP and Trf2 regulate distinct subsets of developmental promoters. A. DNA-affinity purification mass spectrometry enrichment values of TBP and Trf2 across the tested promoter types (pval<0.05), three biological replicates per sample. B. ClustalW alignment of the short and long transcript isoforms of the Drosophila melanogaster TRF2 gene of the C-terminal region from 840-1715 amino acids. Peptides detected from label free mass spectrometric quantification of nuclear lysate from the TRF2-AID cell line are highlighted in yellow. All detected peptides are shared between the two isoforms. C. Noramlize abundance of TRF2 peptides from lab label free mass spectrometric quantification of nuclear lysates form the TRF2-AID cell line. Parental cell line is expressing the Tir1 ligase, while the TRF2-AID cell line is endogenously tagged with 3x-FLAG-AID. 500uM Auxin treatment was performed for 6 hour. D. Western blot of anti-FLAG antibody on the double tagged TBP + Trf2 AID cell line visualizing TBP and Trf2 upon auxin addition, indicating a slower depletion kinetics of the TBP-AID protein. E. Growth curve tracking the number of live cells for 4 days for individual TBP-AID, Trf2-AID, and double TBP+Trf2-AID cell lines. No growth differences are observed upon the individually tagged cell lines, but the double TBP+Trf2-AID cell line shows growth inhibition after addition of auxin. F. PRO-seq signal after TBP or Trf2 depletion (log2 fold change) is plotted for the TBP-dependent genes and Trf2-dependent genes. G. Auxin wash-out experiment in which TBP-AID or Trf2-AID cell lines were treated with auxin for 6 hours and then washed twice and exchanged with fresh medium to remove auxin. qPCR performed on the tested time points on two tested genes indicate they can recover to their original level in the absence of auxin. H. qPCR was performed on an auxin time-course treatment experiment. The tested genes were normalized to Actin5c levels. NLaz was identified from PRO-seq as dependent on TBP but not Trf2, and CG7408 was identified from PRO-seq to be dependent on Trf2 but not TBP. I. In silico LyC and tryptic digestion of the Trf protein reveals predicted detectable peptides, which were not detected in mass spectrometry in our S2 cells, indicating lack of Trf protein expression. J. ClustalW alignment of TBP and Trf. Peptides from TBP detected by mass spectrometry are highlighted in red. Peptides predicted from an *in silico* digest performed on Trf (from panel H) are highlighted in yellow. K. PRO-seq data of individual TBP, Trf2 and double tagged TBP+Trf2 depletion at housekeeping promoters containing DRE, Ohler 1 and Ohler 6 motifs. These promoters are affected only upon depletion of Trf2 and to the same extent upon double depletion, demonstrating that TBP is dispensable for their expression and cannot substitute for Trf2 at these housekeeping promoters.

**Figure S5.**
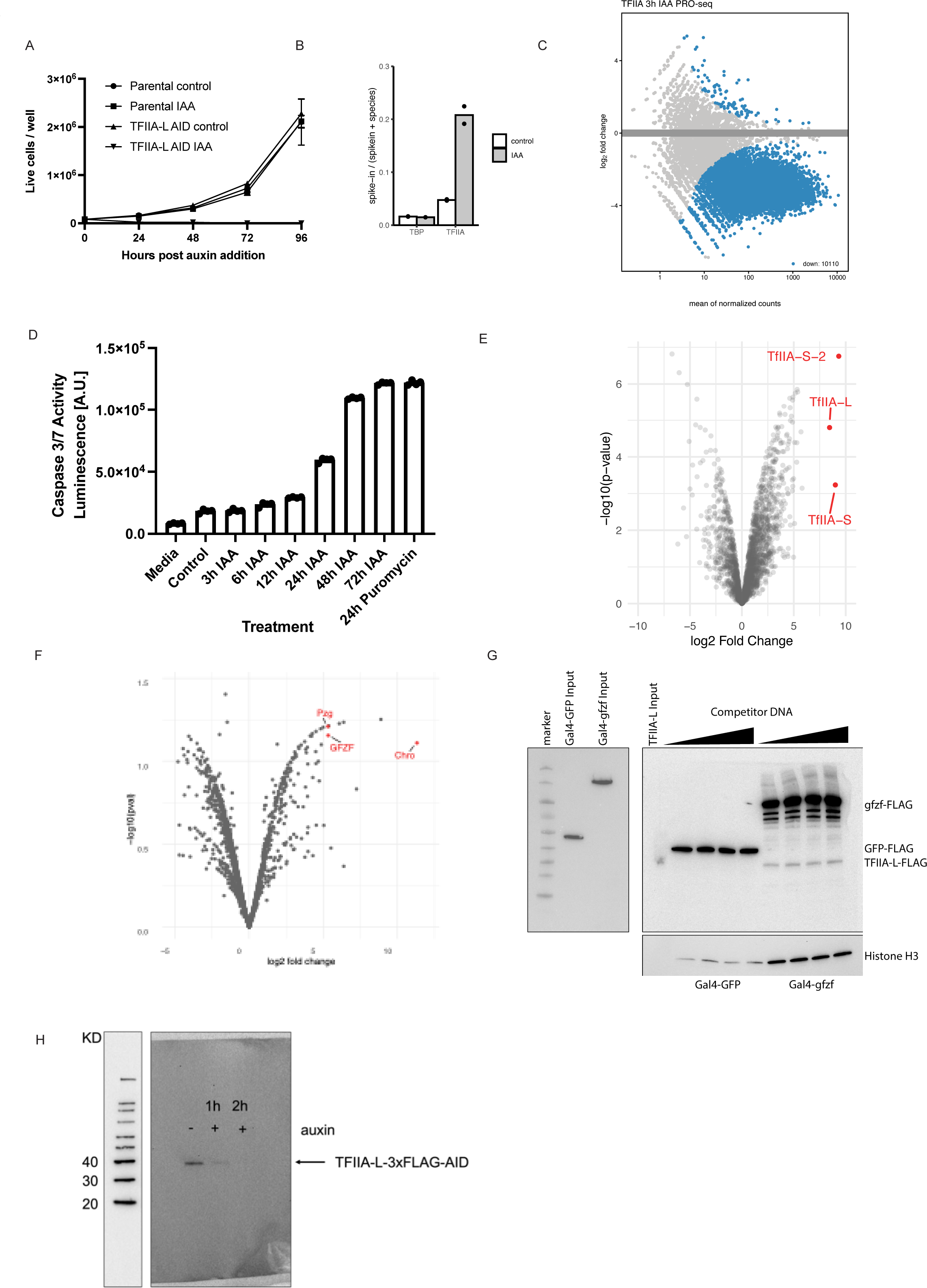
TFIIA is required by all promoters and is recruited by housekeeping cofactors to housekeeping promoters. A. Growth curve of TFIIA-L-3xFLAG-AID cell line and parental Tir1 expressing control over 4 days upon the addition of 500uM auxin. TFIIA-L-AID treated cells die after 24 hours. B. Fraction of reads mapping to the *D. mel* genome (reference species) and the human genome (spike-in) in PRO-seq experiments depleting TBP or TFIIA-L. A ∼4-fold increase in proportion of reads mapping to the spike-in genome is observed only upon depletion of TFIIA-L due to global failure of Pol II transcription in the TFIIA-L-AID cell line. C. MA-plot of PRO-seq data in the TFIIA-L-AID cell line after 3 hours of auxin treatment, showing a global failure of Pol II transcription. D. Caspase 3 and 7 activity was measured with the Promega Caspase 3/7 Glo kit of TFIIA-L-AID cells after addition of auxin at various time points. A positive cell death control was included as a 24 hour treatment of 10ug/ml puromycin. E. Volcano plot of TFIIA-L immunoprecipitation mass spectrometry. TFIIA-L was immunoprecipitated from the endogenously tagged TFIIA-L-3xFLAG-AID cell line using anit-FLAG beads. Enrichment was measured over control immunoprecipitation made from the Tir1 expressing parental cell line which does not contain any FLAG epitope. 3 replicates were performed for each condition. F. Volcano plot of Chromator immunoprecipitation mass spectrometry. Chromator was immunoprecipitated from the Chromator-3xFLAG-AID cell line using an anti-FLAG antibody. Similar Tir1 expressing parental cell line control was used to measure enrichment. Putzig (Pzg) and GFZF are also highlighted. G. DNA-affinity purification assay was performed with a 121bp long housekeeping DRE promoter with 4xUAS sites upstream. Initially, a nuclear extract containing a Gal4-DNA binding domain fusion of GFP or GFZF was incubated with the bead-immobilized promoter DNA (left panel). After the incubation, the extract was removed, and the beads were used for a DNA- affinity purification assay with a nuclear extract containing TFIIA-L-AID-3xFLAG as described in the materials and methods. Sheared salmon sperm DNA was used as competitor DNA at 600ng to 1.6ug per reaction. Elution fractions were ran on an SDS-PAGE gel and blotted with a FLAG antibody (right panel). H. Western blot against FLAG antibody visualizing whole cell lysate from a TFIIA-L c-terminally tagged 3x-FLAG-AID line treated with auxin for two hours. Full degradation of the TFIIA-L beta subunit is visible upon 2 hours of auxin treatment.

**Figure S6.**
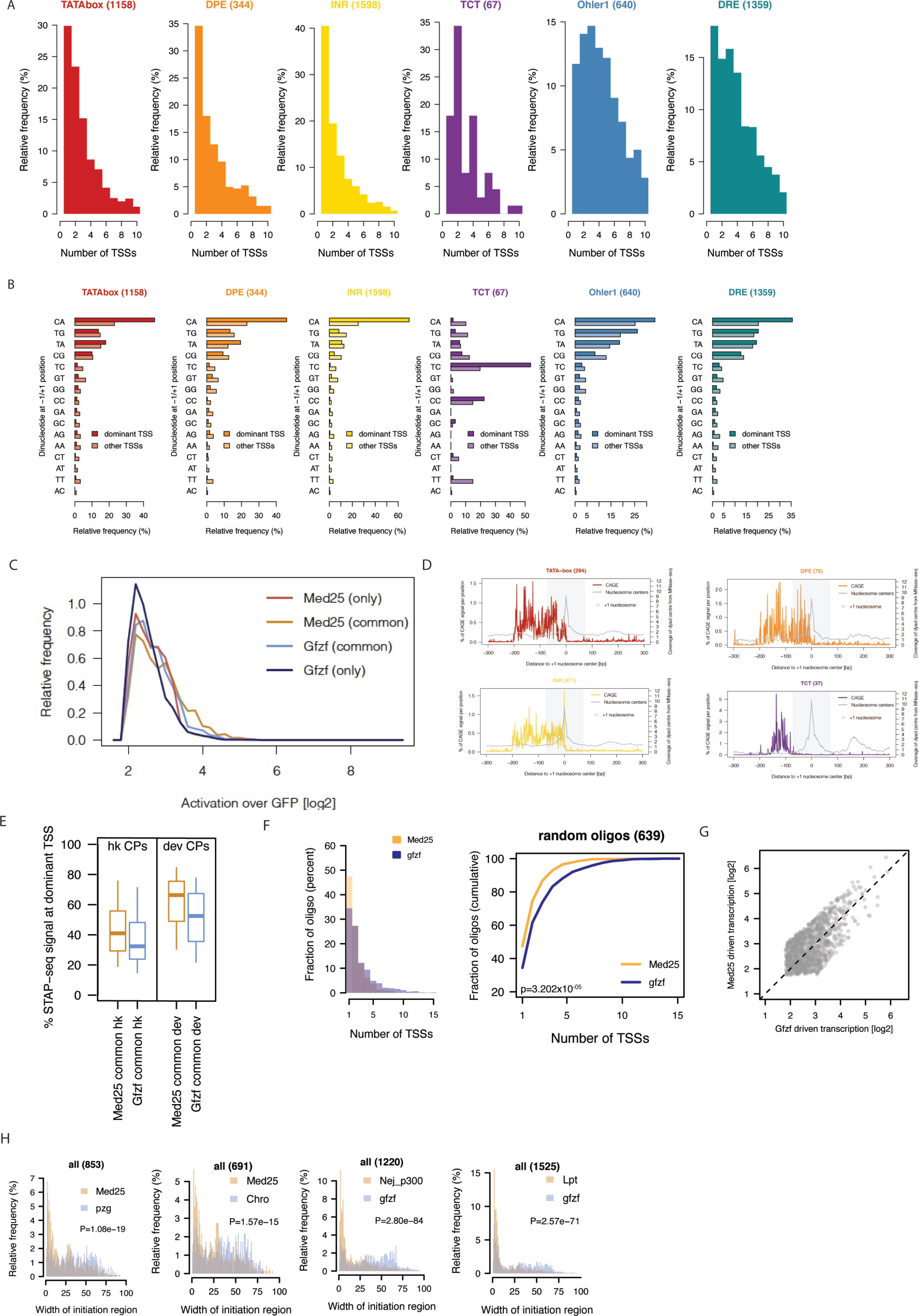
Housekeeping cofactor recruitment is sufficient to recapitulate dispersed transcription initiation patterns. A. The number of CAGE defined TSSs in each promoter type over a 120+/- bp region. TSS was defined as a position having at least 20% CAGE signal as the dominant TSS in the tested region. B. Frequency of dinucleotides at the -1/+1 position for the dominant and secondary TSSs in each promoter type in a 120+/-bp window. C. Fold change (log2) of STAP-seq signal upon GFZF or MED25 recruitment over GFP for oligos that are matched for their activation level by either one of both cofactors. D. Relative CAGE signal per position on all active promoters of the indicated type aligned to the +1 nucleosome centre (point of highest coverage of MNase fragment centres in +1 to +200bp window relative to TSS). E. Percent of STAP-seq signal at the dominant TSS for activation matched oligos (one activated oligo per gene TSS) for housekeeping and developmental promoters that can be activated by both MED25 and GFZF. F. Histogram showing the number of TSSs activated upon GFZF or MED25 recruitment on random regions that are responsive to both cofactors (left). Cumulative plot of the same data (right). P-values: Kolmogorov-Smirnov test. G. Scatter plot of the log2 fold change above GFP (i.e. activation) of promoters by GFZF or MED25 used in the analysis (i.e. matched to be activated to similar extent by both cofactors). H. Histogram representing distribution of the width of the initiation region (i.e. part of the oligo covered by STAP-seq signal) upon recruitment of MED25 or Putzig (Pzg), Med25 or Chro (Chromator), p300 or GFZF, Lpt or GFZF. For each comparison core promoters activated to similar extent by both analyzed cofactors were included. P-values: Wilcoxon rank sum test.

